# Resetting of H3K4me2 during mammalian parental-to-zygote transition

**DOI:** 10.1101/2022.05.05.490401

**Authors:** Chong Wang, Yang Li, Yaqian Wang, Yong Shi, Xiangrui Meng, Wenbo Li, Jia Guo, Kaiyue Hu, Hao Chen, Jiawei Xu

**Affiliations:** The First Affiliated Hospital of Zhengzhou University & Institute of Reproductive Health, Henan Academy of Innovations In Medical Science Zhengzhou, 450000 China; NHC Key Laboratory of Birth Defects Prevention, Zhengzhou, 450000 China; Department of Human Cell Biology and Genetics, School of Medicine; Shenzhen Key Laboratory of Gene Regulation and Systems Biology, Southern University of Science and Technology, Shenzhen, 518000, China

**Author notes:** These authors contribute equally to this work (Chong Wang, Yang Li, Yaqian Wang, Yong Shi).

## Abstract

Upon sperm and oocyte fertilization, the embryo undergoes drastic histone modification reprogramming during pre-implantation development. In this study, we investigated the erasure and re-establishment of H3K4me2 in mouse GV, MII, and embryos using an improved approach called Cleavage Under Targets and Release Using Nuclease (CUT&RUN) with high-throughput sequencing. We found that H3K4me2 extensively occurs as a non-canonical pattern in mouse GV oocytes and early embryos, and parental H3K4me2 was erased by maternal LSD2 during forming the pronucleus in the zygote. The H3K4me2 was erased from GV to MII oocyte and was re-established in the late two-cell stage, removing this epigenetic burden is crucial for ZGA. We then revealed the H3K4me2 locates widespread in the CpG-rich and hypomethylated regulatory regions in the four-cell stage, and the eight-cell stage, but dramatic changes occurred in the inner cell mass (ICM) in mouse embryos. Then, these CpG-rich H3K4me2 regulatory regions became either activated or repressed. To summarize, in this study, we elucidated the pattern of H3K4me2 transition from parent to zygote and H3K4me2 profile during early embryo development. Our findings might provide deeper insights into epigenetic reprogramming during early development and in vitro fertilization in mammalian.

**Highlights:**

1. H3K4me2 showed differential patterns in sperms and oocytes; non-canonical pattern occurred in GV oocytes, but canonical patterns were found in the sperm cells of mice.
2. The H3K4me2 peaks of GV were almost erased in MII oocytes and reconstructed after the major ZGA occurred. Maternal LSD2 was found to regulate H3K4me2 resetting during the parental-to-zygote transition. H3K4me2 may impose an “Epigenetic Burden” on ZGA.
3. Non-canonical H3K4me2 pattern was found in mammalian embryos, however, pervasive H3K4me2 in CpG-rich regulatory regions were found in four-cell and eight-cell stage embryos; The CpG-rich H3K4me2 regulatory regions were either activated or repressed in the ICM.

## INTRODUCTION

Drastic epigenetic reprogramming occurs during early mammalian embryo development. Epigenetic marks, such as DNA methylation and histone modification, play important roles in the epigenome reprogramming process^1,2^. Epigenome reprogramming during the pre-implantation embryo development stage erases the epigenetic memory of oocytes and sperm cells and allows the zygote to establish a new epigenetic landscape, which is relevant for early development and is essential for maternal zygotic transition (MZT) and differentiation^1,3^. Deficiencies in the reconstruction of the epigenome can result in developmental defects, such as embryonic arrest^3^. Histone modifications are fundamental epigenetic regulators that control gene expression and many key cellular processes^4-6^. Different types of histone modifications can occur in specific genomic regions and perform distinct functions^3^. Most histone modifications occur around transcription start sites (TSSs) in the embryos; this feature is shared by all somatic cells^7-9^. A similar pattern of histone modification commonly occurs in many species and cell lines^10-12^, and thus, this distribution pattern is known as the ‘canonical’ pattern^13^. Highly pervasive H3K27me3 is found in regions depleted of transcription and DNA methylation in oocytes, whereas extensive loss of promoter H3K27me3 occurs at *Hox* and other developmental genes after fertilization, accompanied by global erasure of H3K27me3 from sperm but inheritance of distal H3K27me3 from oocytes; both H3K4me3/H3K27me3 bivalent promoter marks are restored at developmental genes^14^.

The K-residues at the N-terminus of histone H3 undergo mono-(H3K4me1), di-(H3K4me2), or tri-methylation (H3K4me3), catalyzed by a family of proteins that contain a SET domain; the methylation of a different number of methyl groups added represents different chromatin status^15-17^. Different numbers and kinds of histone modification marks are associated with different gene expression states^18^. H3K4me3 is usually enriched near transcription start sites (TSS) that are associated with the activation of nearby genes^19^. H3K4me1 is a global marker of enhancers; regions enriched with H3K4me1 and H3K27ac are identified as super enhancers^20-22^. H3K4me1/H3K4me2/H3K4me3 is enriched on both sides of the TSS and plays diverse and important roles in regulating gene expression in mammals^23,24^. Some studies found that the H3K4me2 distribution pattern is like the H3K4me3 distribution pattern in plants^25,26^. Like LSD1, LSD2 (KDM1B) belongs to the FAD-dependent amine oxidase family and shares similar substrate specificity with LSD1, which can function as a H3K4me2 demethylase independently of LSD1^27-29^. The mechanism underlying H3K4me2 reprogramming in the early development of mammals is still unknown, and the importance of the inheritance of the H3K4me2 programming from parents to offspring needs to be determined.

H3K4me2 can coordinate the recovery of protein biosynthesis and homeostasis following DNA damage, and it is required for the growth and survival of organisms^30^. H3K4me2 enrichment in plants is negatively correlated with the gene transcript level. H3K4me2 in rice and arabidopsis specifically distributes over the gene body region and functions as a repressive epigenetic mark^31^. H3K4me2 was found to be a key epigenetic factor in the chicken PGC cell lineage determination, and a narrow domain activated PGC-specific expression genes and signaling pathways^32^. H3K4me2 can occur as early as the embryonic stem cell stage, and its distribution is dynamic and progressive throughout the mouse brain differentiation and tissue development stages^33^. About 90% of the transcription factor binding regions (TFBRs) overlap with the H3K4me2-enriched regions, and along with the H3K27ac-enriched regions, they can greatly reduce false positive predictions of the TFBRs. Additionally, the stronger signals of H3K4me2 enrichment are more likely to overlap with the TFBRs than with weaker signals^34^. A subgroup of genes enriched with high levels of H3K4me2 was found within their gene body linked to T cell functions, which was different from the TSS-centered enrichment of housekeeping genes^35^. Many regions of the sperm epigenome bearing H3K4me2 are present in spermatogonia, suggesting an early establishment of H3K4me2 in spermatogenesis, which is a highly conserved process^36^.

In this study, we elucidated the mechanism underlying H3K4me2 reprogramming during the early development of mammals. We optimized the Cleavage Under Target and Release Using Nuclease (CUT&RUN)^37,38^ methods using H3K4me2 antibody to as few as 50 cells and revealed unique H3K4me2 modification patterns and epigenetic reprogramming that occur during the parental-to-zygote transition in mice. We also showed that H3K4me2 resetting depends on maternal LSD2 and parental H3K4me2 acts as an “Epigenetic Burden” on mammalian ZGA.

## Results

### Genome-wide profiling of H3K4me2 in mouse oocytes and early embryos

The epigenome underwent significant reprogramming during early development. To determine how H3K4me2 is inherited from the sperm and oocyte by the zygote during early development, we investigated H3K4me2 in mouse gametes and pre-implantation embryos using CUT&RUN methods. We optimized the CUT&RUN protocol (CUT&RUN ; Methods) using H3K4me2 antibody. We also acquired high-quality data using only 50 cells. Most peaks could be retrieved using only 50 cells (Figures S1A and S1B). Then, we collected GV oocytes, MII, PN5, early two-cell embryos, late two-cell embryos, four-cell embryos, eight-cell embryos, and the inner cell mass (ICM) of mice. Mouse embryonic stem cells (mESC) were used as control. We profiled H3K4me2 in oocytes and early embryos using CUT&RUN and immunofluorescence (IF) techniques. The results of all replicates were highly consistent (Figure 1A; Figures S1A, B, and C). We identified 57,455, 25,073, 110,517, 126,318, 62,575, and 69,154 H3K4me2 peaks in the GV oocytes, late two-cell embryos, four-cell embryos, eight-cell embryos, ICM, and mESC, respectively. Most of the H3K4me2 was erased from GV to MII during oocyte development. The paternal H3K4me2 was also erased after fertilization, and it was re-established in the late two-cell stage (Figures 1A-B). We also demonstrated the H3K4me2, ATAC-seq, and mRNA expression of several developmental key genes (Figure 1C). We found that H3K4me2 positively correlated with H3K4me3 and chromatin accessibility in early embryos (Figures S1D-E). The H3K4me2 genome-wide pattern of four-cell embryos showed a strong association with eight-cell embryos: the H3K4me2 of four-cell and eight-cell embryos was in one cluster, and the correlation was high (R = 0.93) (Figures S1F-G).

**Figure 1.**
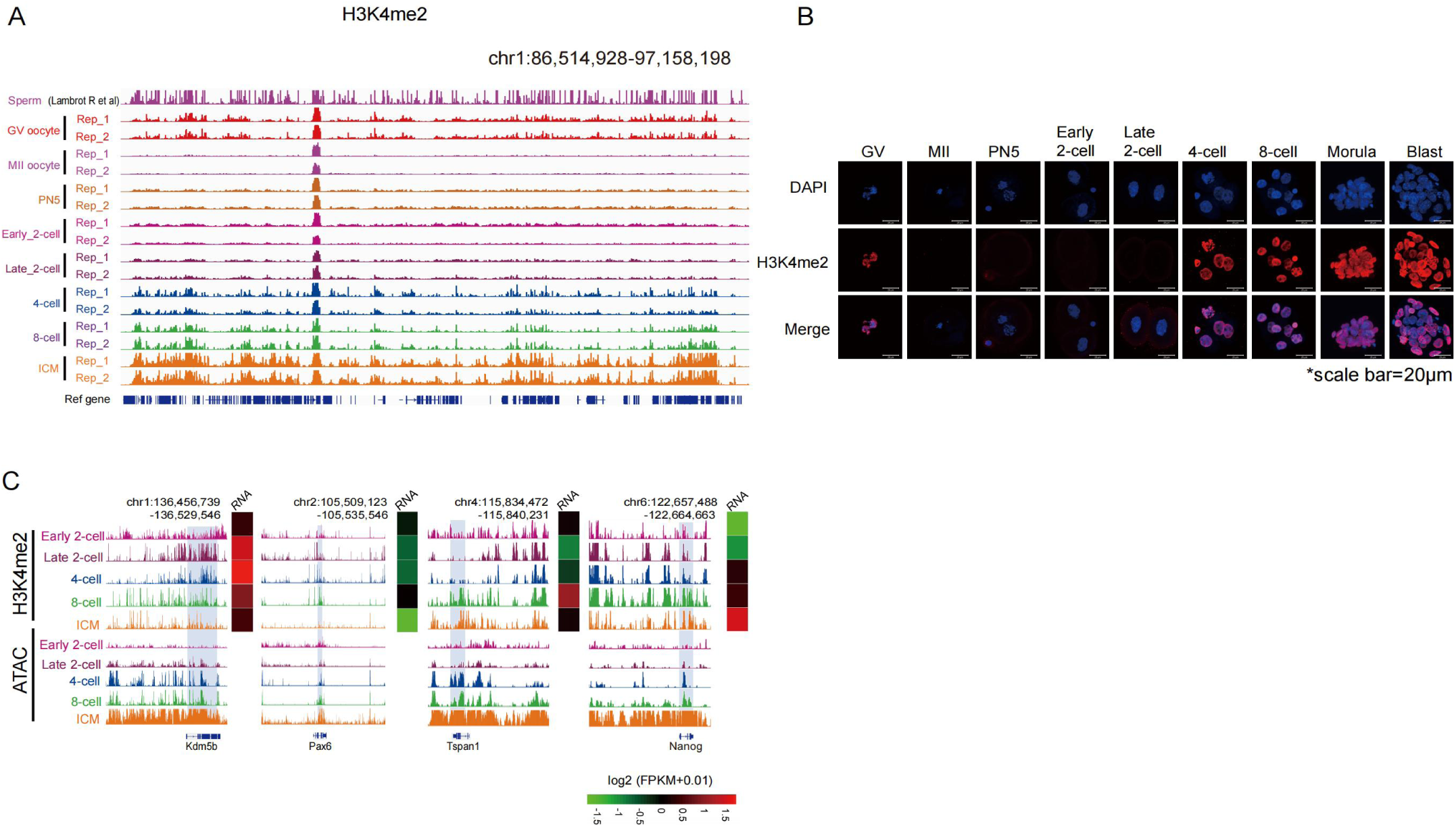
Mapping H3K4me2 in mouse (C57BL6/N) gametes and pre-implantation embryos. (A) The IGV view shows the global landscape of H3K4me2 signals in mouse sperm, GV oocytes, MII oocytes, and pre-implantation embryos. The H3K4me2 data in sperm was obtained from ENCODE (Lambrot et al.). ICM was isolated from the blastocyst. (B) Immunofluorescence of H3K4me2 in mouse (C57BL6/N) GV oocytes, MII oocytes, PN5 zygote, early two-cell embryo, late two-cell embryo, four-cell embryo, eight-cell embryo, morula, and blastocyst; the scale bar is also shown. (C) H3K4me2 enrichment near typical genes and their expression levels are shown. H3K4me2 signal, ATAC-seq, and gene expression are shown.

### Distinct H3K4me2 pattern between sperm and oocyte

We systematically characterized the epigenome of mouse gametes. In the analysis, we included published data on DNA methylation^39^, H3K4me3^40^, and ATAC-seq of early embryos^41^, as well as the H3K4me2 data of sperm^42^. Non-canonical patterns of H3K4me3, enriched broad domains in promoters and distal regions, are present in mouse oocytes as broad domains in partially methylated domains (PMDs)^43,44^. H3K4me2 shows a non-canonical pattern in the GV oocytes and sperm of mice (Figures 2A-B and E-F). Mature sperm and oocytes are in a state of inactive transcription^45,46^; however, strong ChIP-seq and CUT&RUN signal enrichment of H3K4me2 was found to be related to transcription^47^. These findings suggested that H3K4me2 might be deposited during gametogenesis and retained in mature gametes. To confirm this speculation, we performed a K-means cluster analysis for H3K4me2 in gametes. By comparing promoter H3K4me2 at the sperm and GV stages, we found that the promoters could be divided into four groups (Figure 2A). The promoters in the first group (sperm-specific H3K4me2) were involved in fertilization and germ cell development, and these markers were erased in the later stages (Figures 2A and 2C). In contrast, the promoters in the second group (GV-specific H3K4me2) were associated with mitosis and gamete generation. They played a role in GV generation during oocyte development and were re-established in the late stages (Figures 2A and 2C). The promoters in the third group (sperm-GV share H3K4me2) were related to mRNA processing and translation. The signals of these regions were erased (MII, zygote, and early two-cell stage) and reconstructed (at the late two-cell stage) during development (Figures 2A and 2C). The promoters in the fourth group had a low H3K4me2 level at all stages (Figure 2A).

**Figure 2.**
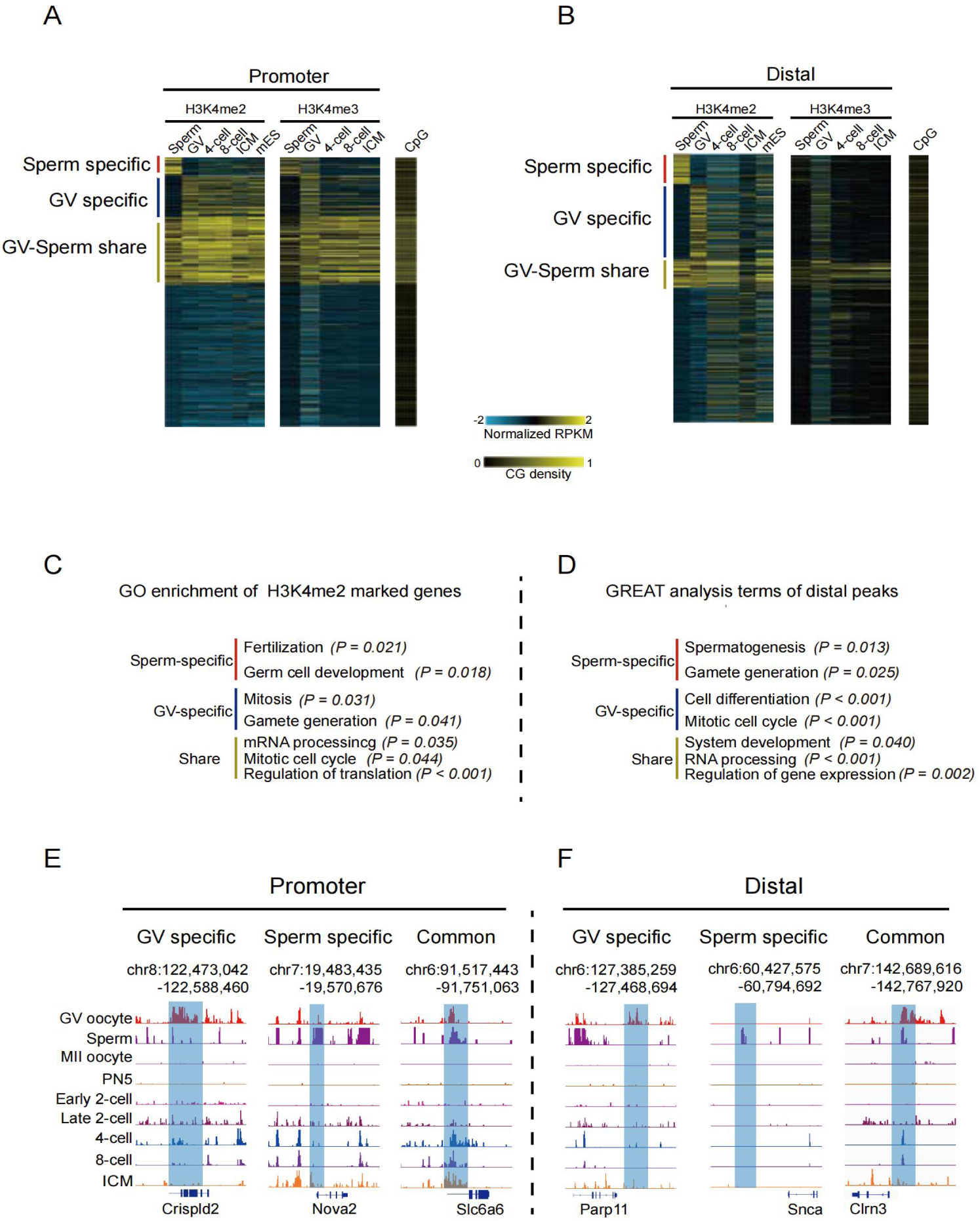
Comparison of H3K4me2 between sperms and oocytes. (A-B) Heatmaps showing H3K4me2 and H3K4me3 expressed genes between sperm and GV oocyte in the promoter and distal region at different stages. CpG densities are also mapped and shown. (C) The enriched GO terms of H3K4me2 marked genes in GV and sperm-specific or shared genes are shown along with the associated p-values. (D) The results of the GREAT analysis for distal peaks in GV and sperm-specific or shared genes are shown along with the associated p-values. (E-F) The IGV views show H3K4me2 expressions in GV and sperm-specific or shared genes at the representative promoters (left, shaded) or distal accessible regions (right, shaded).

The H3K4me2 enrichment in these promoters was consistent with the changing dynamics of H3K4me3 (Figure 2A). Specifically, promoters with high CpG density were preferentially marked by H3K4me2 in the sperm-GV share group (Figure 2A). Next, we investigated H3K4me2-marked distal regions in gametes. Sperm-specific groups are involved in gamete generation (Figure 2D). GV-specific distal H3K4me2 showed functions in cell differentiation and mitotic cell cycle (Figure 2D). Both were erased after fertilization and not reconstructed during embryo development (Figure 2B). H3K4me3 signals did not occupy the sperm-specific/GV-specific H3K4me2 distal regions (Figure 2B). The GV-sperm-shared distal H3K4me2 regions were also marked by H3K4me3. Those genes participated in system development, RNA processing, and regulation of gene expression (Figures 2B and 2D). Compared to GV, sperm H3K4me2 was enriched in a fully methylated promoter, but the distal H3K4me2 of both sperm and GV were enriched in the hypomethylated region (Figures S2A-D). These results indicated that differential H3K4me2 modification patterns occurred between sperms and oocytes.

### Resetting H3K4me2 in pre-implanted embryos

Significant epigenetic reprograming and chromosome remodeling associated with H3K4me3, H3K27me3, H3K36me3, and H3K27ac occurs during early mammalian development^44,48,49^. In another study, we found that epigenome reboot occurred during the early developmental stage in humans^50^. We investigated how parental histone mark H3K4me2 at regulatory elements are reprogrammed after fertilization. We found that GV and sperm H3K4me2 peaks were almost erased in PN5 and re-established along with zygotic genome activation (ZGA). This result was further confirmed by IF analysis, which showed that the reconstruction of H3K4me2 might start from the late two-cell stage (Figures 1B and 3B). It is worth noting that previous studies using different immunofluorescence parameters reported detectable H3K4me2 signals at the zygote and two-cell stages^51^. This discrepancy may arise from differences in confocal microscope settings, antibody sensitivity, or sample processing. In our study, we used consistent parameters across all stages from GV to blastocyst; when we specifically optimized the acquisition parameters for zygotes and two-cell embryos, weak signals could be observed. The H3K4me2 demethylases KDM1B (LSD2) were also highly expressed in growing oocytes but showed decreased expression in MII oocytes (Figure S3A, see also revised heatmap). We have re-analyzed the expression data and corrected the heatmap normalization; the revised Figure S3A now accurately shows that Kdm1b is highly expressed in growing oocytes, with lower levels in 8-week and MII oocytes, consistent with its role in maternal imprint establishment. The CUT&RUN data, which showed small H3K4me2 peaks (related to other stages), reappeared at the late two-cell stage (Figure 3A). We concluded that the IF analysis could not detect the newly established weak H3K4me2 signal in the late two-cell stage. H3K4me2 was correlated with the activation of Prdm9 and Kmt2c at the four-cell stage (Figure S3A), and the signal was strong enough to be detected at the four-cell and eight-cell stages (Figures 3A-B) using both the CUT&RUN and IF techniques. The four/eight-cell H3K4me2 targets only partially overlapped with those in sperm or oocytes, and these overlapped targets were mainly enriched in development-associated pathways or processes (Figure 3C). Additionally, gamete-specific H3K4me2 targets were involved in the development and generation of gametes, whereas four/eight cell-specific targets were associated with cell communication and metabolic processes (Figure 3C). These findings indicated that the resetting of H3K4me2 involves distinct sets of genes that are closely related to the development of embryos from mouse gametes to early embryos. About 1% of the four/eight cell H3K4me2 occurred in the hypomethylated regions as early as the late two-cell stage (Figures S3B-D). These results indicated that the resetting of H3K4me2 starts from the late two-cell stage but is mostly re-established at the four/eight-cell stage.

**Figure 3.**
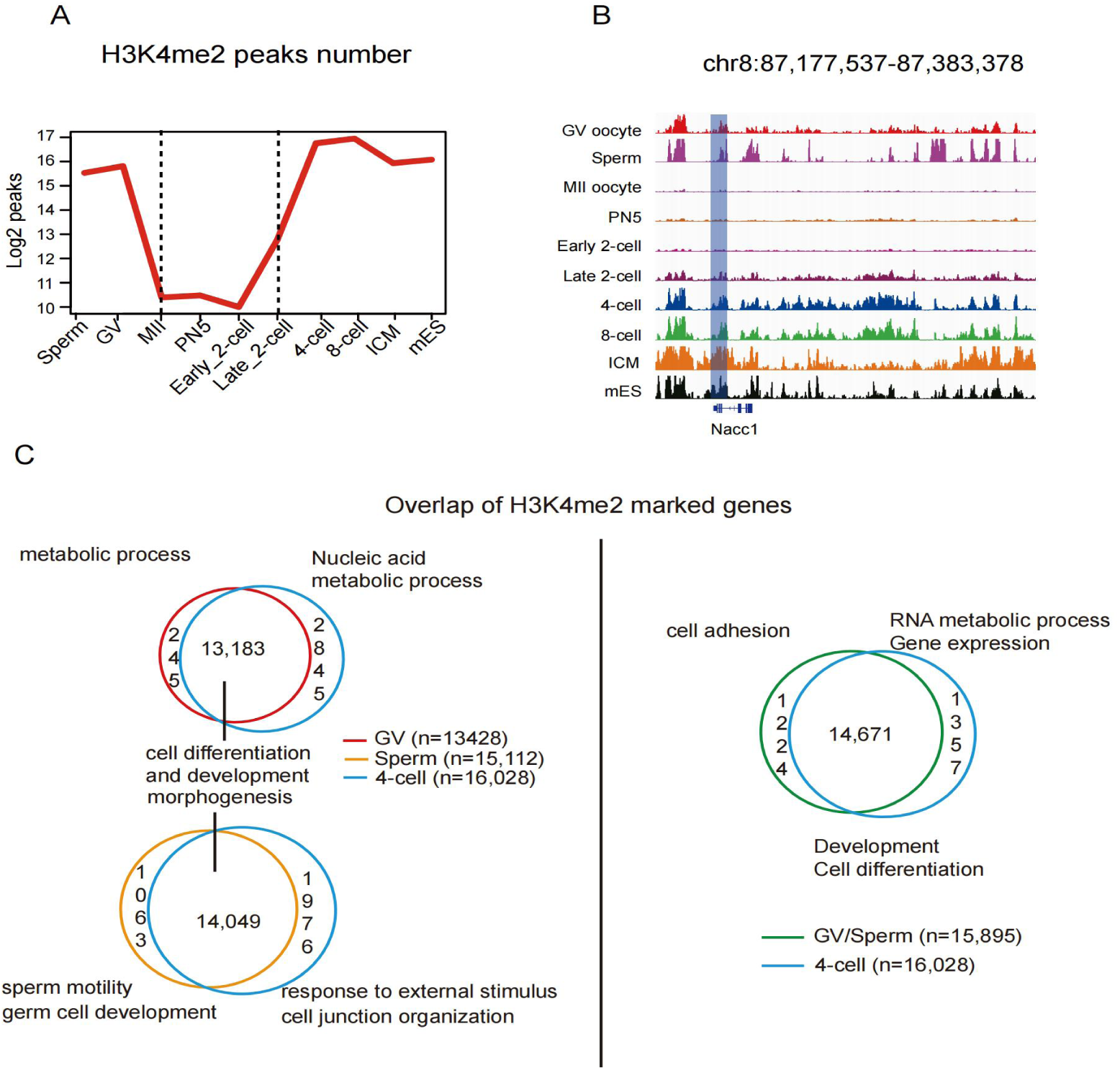
Resetting of H3K4me2 during mouse pre-implantation development. (A) The number of H3K4me2 peaks in mouse embryo stem cells (mESCs), mouse gametes, and pre-implantation embryos was calculated. (B) The IGV view shows H3K4me2 signals in mESCs, mouse gametes, and pre-implantation embryos; Ncc1 gene sites are also shown. (C) Venn diagrams illustrate the overlap of H3K4me2 marked genes between sperm/GV oocytes, separately (left) or jointly (right), and four-cell embryos. The enriched GO terms for the shared and non-overlapping genes are also shown.

### The inhibition of LSD2 increased the expression of H3K4me2, which impaired the development of embryos in vitro

Histone post-translational modifications are coordinated by histone methyltransferases and demethylases^52^. KDM1B, also known as LSD2, is the only demethylase homologous to LSD1 in mammals^29^. Although they share the H3K4me1/2 demethylase function, our analysis showed that Kdm1a was rarely expressed during mouse embryonic development (Figure 4A). While, LSD2, which is required to establish the maternal imprinted gene^53^, was highly expressed before the early two-cell stage and decreased after the late two-cell stage (Figures 4A and S3A).

**Figure 4.**
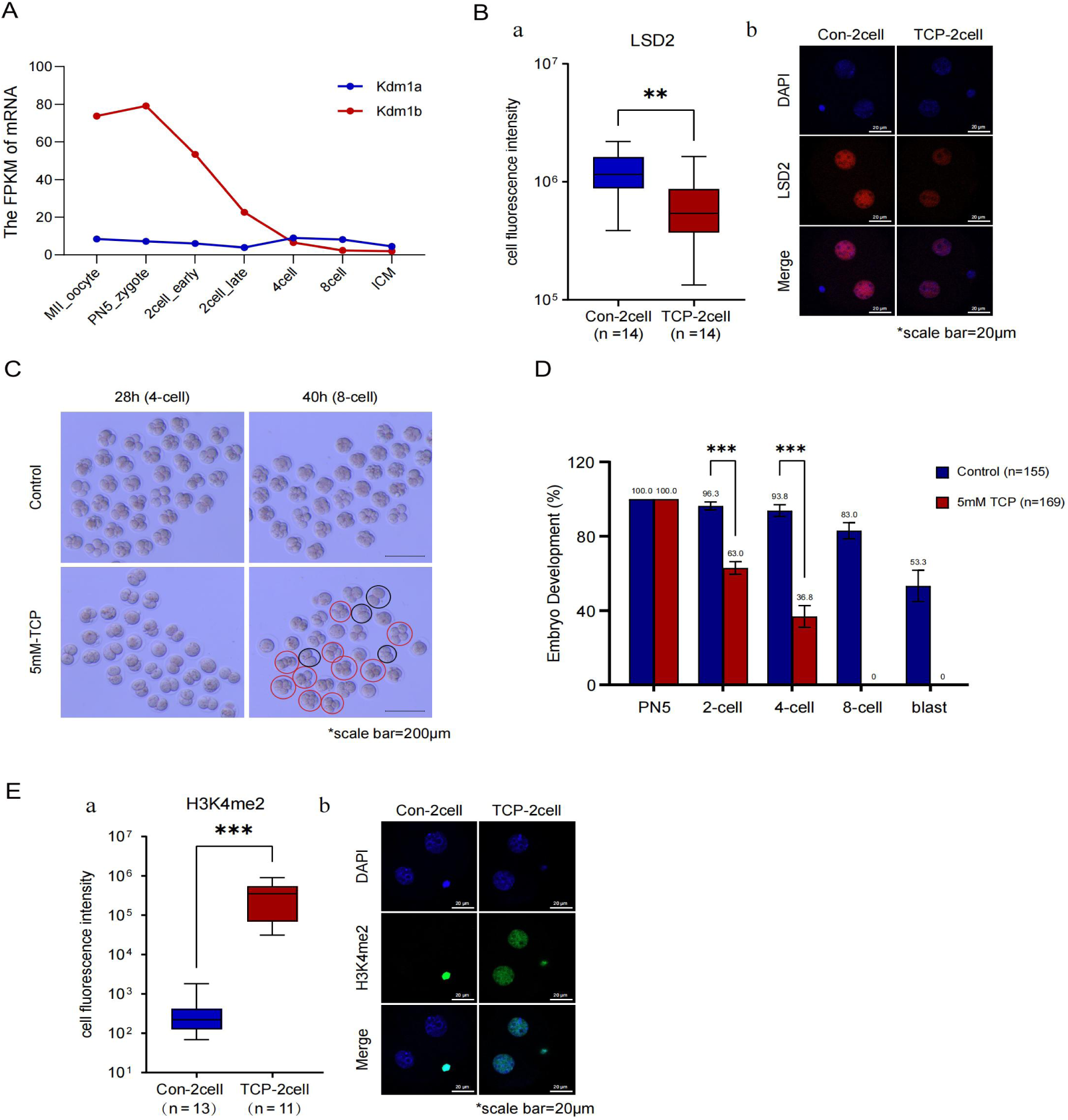
The inhibition of LSD2 resulted in abnormal embryonic development. (A) The dynamic changes in the Kdm1a (blue line) and Kdm1b (red line) levels in oocytes and early embryos (based on published RNA-seq data43). (B) a.The immunofluorescence intensity of LSD2 in the experimental group (5 mM TCP) and the control group are shown using a box diagram. b. Immunofluorescence images showed the immunofluorescence intensity of LSD2 of two-cell embryos in the experimental group (5 mM TCP) (right) and the control group (left); the scale bar is also shown. (C) The developmental morphology of mouse embryos in the experimental group (5 mM TCP) and the control group after 28 h and 40 h of culture are shown along with the scale bar. (D) The figure illustrates the embryonic development of mice at different stages after culture in vitro in the experimental group (5 mM TCP) and the control group. (E) a.The immunofluorescence intensity of H3K4me2 in the experimental group (5 mM TCP) and the control group are shown using a box diagram. b. Immunofluorescence images showed the immunofluorescence intensity of H3K4me2 of two-cell embryos in the experimental group (5 mM TCP) (right) and the control group (left); the scale bar is also shown.

Tranylcypromine (TCP) can inhibit the expression of LSD2^54^. To investigate the effect of the expression of H3K4me2 on embryonic development, mouse PN5 embryos were cultured in vitro with three different TCP treatment groups (3.6 mM, 5 mM, and 10 mM) and a control group. The inhibition in the presence of 3.6 mM TCP was not prominent, and some embryos could develop to the eight-cell stage at 65 h. When 10 mM TCP was present, embryonic development was strongly inhibited, and the embryo could not develop to the two-cell stage (Figure S4A). When 5 mM TCP was present, the embryos could pass through the two-cell stage with a significant developmental delay. Some embryos developed till the four-cell stage, and then, were completely arrested. These results showed that TCP affected embryonic development in a dose-dependent manner and caused developmental arrest. As no prior study has reported the precise effective concentration of TCP for inhibiting LSD2 in mouse embryos, our dose-response experiment empirically determined that 5 mM TCP was sufficient to cause significant H3K4me2 accumulation and developmental arrest without immediate lethality (see Methods).We concluded that the TCP concentration of 5 mM in the experimental group (5 mM TCP group) could be used to determine whether the abnormal expression of H3K4me2 affects the development of embryonic cells, especially the occurrence of ZGA.

We found that adding TCP to culture embryos significantly reduced the expression of LSD2 in embryos (Figure 4B). After 28 h of culture, some of the embryos in the Control group developed to the four-cell stage, while those in the 5 mM TCP group remained at the two-cell stage (Figures 4C and S4B). The development of the embryos was significantly delayed after TCP was added compared to the time taken for the embryos in the Control group to develop. The number of embryos at each stage in the 5 mM TCP group (n = 169) and the Control group (n = 155) was counted and statistically analyzed (Figure 4D). The results showed that 62.96% of embryos in the 5 mM TCP group developed to the two-cell stage (Control: 96.29%), and 36.82% of embryos developed to the four-cell stage (Control: 93.83%). All embryos in the TCP group were arrested at the four-cell stage (*P*< 0.001).

To test whether LSD2 specifically functions as H3K4me2 demethylase in mouse embryos, immunofluorescence staining of cells in the 5 mM TCP group and the Control group was performed using H3K4me1, H3K4me2, and H3K4me3 antibodies, respectively. We calculated the immunofluorescence intensity of individual embryos and found no significant difference in the changes in the expression of H3K4me1 between the 5 mM TCP group and the Control group (Figure S4C); also, the changes in the expression of H3K4me3 was not statistically significant (Figures S4C). In contrast, when LSD2 expression was inhibited by TCP, the expression of H3K4me2 at the two-cell stage was significantly higher than the expression of H3K4me2 in the cells of the Control group (*P*< 0.001) (Figure 4E).Elimination of the demethylase action of LSD2 increased the expression of H3K4me2, which delayed and arrested embryonic development. We also found that LSD2 specifically demethylated H3K4me2 in mammalian embryos.

### Parental H3K4me2 was erased by maternal LSD2 during forming the pronucleus in the zygote

During spermatogenesis, most of the histone modifications are replaced by protamine, but a few histones, including H3K4me2, remain conserved^55^. During the early stages of zygote formation, female and male pronuclei are transcriptionally silent, and the development process is regulated by maternal factors. In our study, during the early stage of zygote formation, the expression of H3K4me2 in PN5 was lower than that in PN3 (*P* < 0.001) (Figures 5A-B), which indicated that H3K4me2 signaling was gradually reduced during the fusion of sperm and egg. When PN3 embryos were cultured to the PN5 stage under 5 mM TCP conditions, the fluorescence signal of H3K4me2 increased significantly (*P* < 0.001) (Figures 5A-B), suggesting that the LSD2 might response the removal of parental H3K4me2 during normal embryonic development. The immunofluorescence values of H3K4me2 in female and male pronuclei were assessed to verify whether the effect of LSD2 inhibition on H3K4me2 was different between male and female pronuclei. When LSD2 was inhibited, the expression of H3K4me2 increased significantly in female and male pronuclei, especially in the male pronucleus (334% increase in the male pronucleus and 155% increase in the female pronucleus) (Figure 5C). Sperm carrying H3K4me2 might be more affected by LSD2 demethylase, creating a low H3K4me2 expression environment for zygotic genome activation. To summarize, parental H3K4me2 was erased by maternal LSD2, the H3K4me2 in the male pronucleus was erased faster than that in the female pronucleus, and maternal LSD2 specifically demethylated H3K4me2.

**Figure 5.**
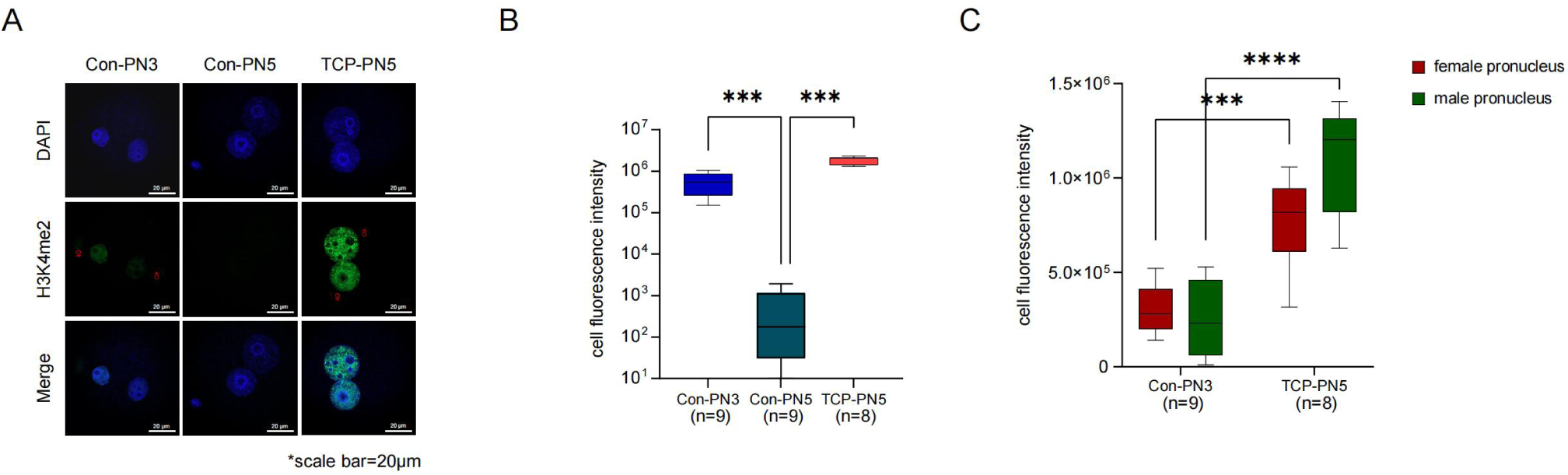
Parental H3K4me2 was erased by maternal LSD2 during sperm-egg fusion. (A) Immunofluorescence images showed the immunofluorescence intensity of H3K4me2 in single-cell embryos in the experimental group (5 mM TCP) and the control group (C); the scale bar is also shown. (B) The immunofluorescence intensity of H3K4me2 in single-cell embryos in the experimental group (5 mM TCP) and the control group are represented as box and whisker plots. (C) The immunofluorescence intensity of H3K4me2 in male and female embryonic prokaryotes of the experimental group (5 mM TCP) and the control group are represented as box and whisker plots.

### H3K4me2 participated in the mammalian ZGA and chromatin accessibility regulation

We used the CUT&RUN technology to detect H3K4me2 throughout the genome and performed gene annotation of the differential expression peaks of H3K4me2 in the TCP-treated group and the control group. We found that the H3K4me2 signal increased in 251 genes and decreased in 194 genes (Fold changes > 2, *P* < 0.01) (Figure 6A). The enrichment analysis of the genes from which H3K4me2 differential peaks originated showed that the molecular function was mainly enriched in the transcription regulation of the promoter of RNA polymerase Ⅱ, and 35 differential genes were involved in this regulation (Figure 6B).

**Figure 6.**
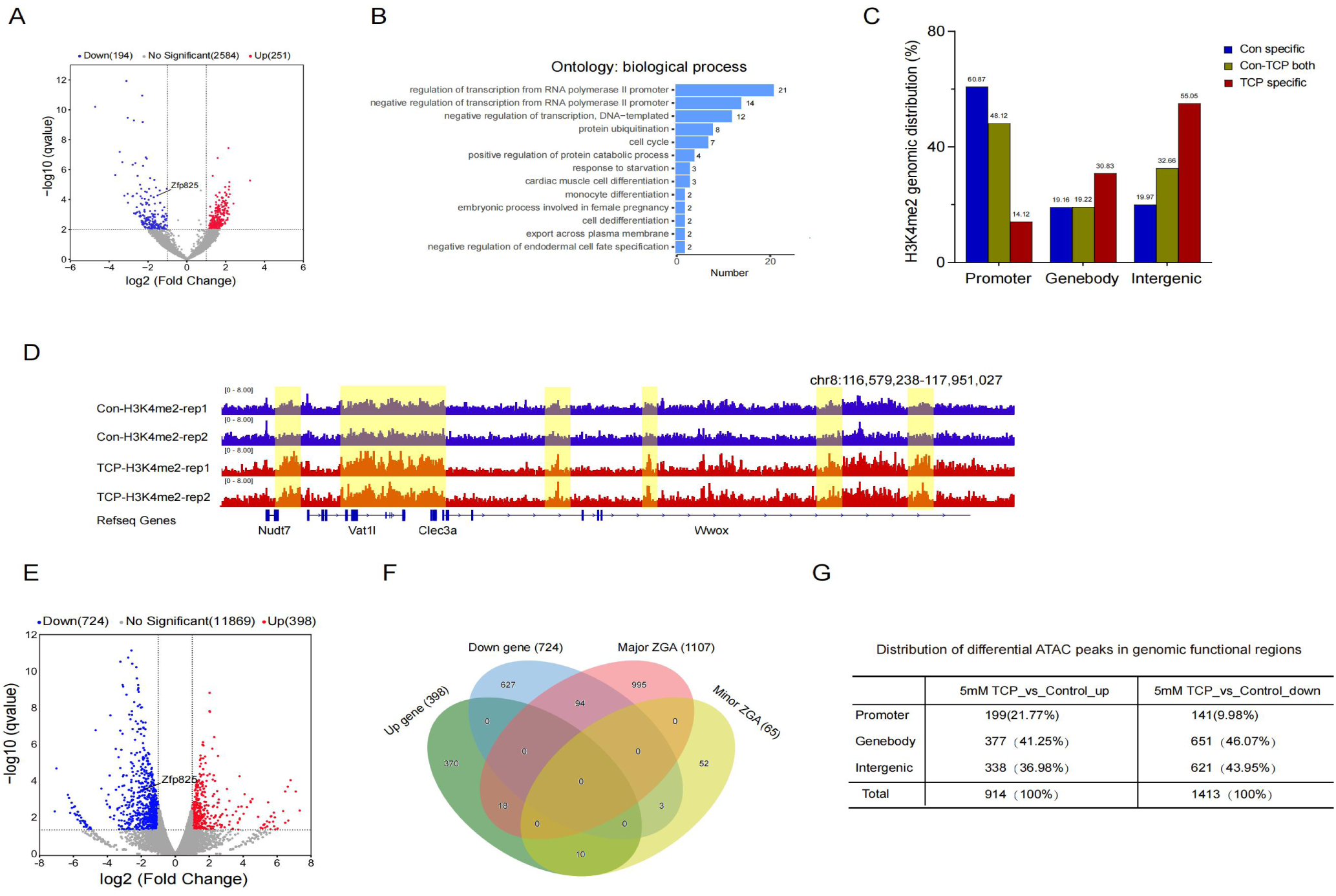
Inhibition of LSD2 affects gene expression. (A) A volcano map of genes with differential H3K4me2 expression after TCP inhibition of LSD2 expression. (B) The GO analysis showed enrichment of biological processes after an increase in the expression of H3K4me2. (C) The peak distribution area of H3K4me2 in the experimental group (5 mM TCP) and the control group (D) The IGV view shows the H3K4me2 signals in typical genes in the experimental group (5 mM TCP) and the control group. (E) A volcano map of the differential expression of H3K4me2 genes in the experimental group (5 mM TCP) and the control group. (F) The Venn diagram illustrates the intersection of differentially expressed genes and minor ZGA and major ZGA genes between the experimental group (5 mM TCP) and the control group (G) The distribution of differential ATAC peaks in the functional regions of the genome.

Unlike LSD1, which binds to the promoter region, LSD2 is absent in the promoter region and enriched in the intronic, exonic, and intergenic regions^29^. The distribution of H3K4me2 changed when LSD2 expression was inhibited (Figures 6C-D). The increase in the H3K4me2 signal mainly occurred in the intergenic and genebody regions (intergenic region: 55.05%, genebody region: 32.66%). We speculated that H3K4me2 located in these regions might be erased by LSD2. However, the deletion of LSD2 did not increase the H3K4me2 signal in the promoter region.

Histone modifications can regulate early embryonic development in mammals by affecting the expression of developmental genes and reprogramming the chromatin structure^1,3,9^. RNA-seq data of mouse germ cells and embryos (refer to GEO database: GSE71434) were used to screen minor/major ZGA genes. We examined the gene expression levels of two-cell embryos in the 5 mM TCP group and the Control group (fold changes > 2, *P*< 0.05) and found that 398 genes were up-regulated and 724 genes were down-regulated in the two-cell stage embryos of mice after LSD2 was inhibited (Figures 6E-F). Among these differentially expressed genes, 94 down-regulated genes were major ZGA genes, and 18 up-regulated genes were major ZGA genes. The Zfp825 gene played a positive regulatory role in RNA polymerase II promoter transcription^34^. Inhibition of LSD2 can lead to a reduction in the distribution of H3K4me2 on the Zfp825 gene and a weakening of the expression of Zfp825. These changes may affect the frequency, rate, or extent of RNA polymerase II promoter transcription.

An increase in the expression of H3K4me2 changed the accessibility of chromatin, of which 1,413 peaks were up-regulation and 914 peaks were down-regulated (fold change > 2; *P*< 0.05) (Figure 6G). Peak-related gene screening of the genes near the promoter region (TSS ±2.5 kb) showed that 240 peak-related genes were up-regulated and 197 peak-related genes were down-regulated. The transcription of genes requires the opening of the tight structure of chromatin to allow the binding of regulatory factors. The up-regulation of H3K4me2 was associated with altered chromatin accessibility in some developmental genes, which in turn could affect developmental progression. The inhibition of LSD2 expression leads to changes in the distribution of H3K4me2 on genes, abnormal expression of genes associated with ZGA, and disorders related to chromatin opening; these changes together may inhibit normal embryonic development.

### Non-canonical H3K4me2 in mammalian embryos and pervasive H3K4me2 in CpG-rich regulatory regions in the four-cell and eight-cell embryos of mice

In GV oocytes and post-ZGA stages, H3K4me2 was enriched in both canonical gene promoters and partially methylated domains (PMDs) (Figures 7A-D). Our results showed that H3K4me2 might be re-established in the late two-cell stage, and strong promoter H3K4me2 enrichment might be completed in the four-cell stage and maintained until the eight-cell stage. We also found that the H3K4me2 signals occupied broader regions than those at other stages (Figure 1A). About 65% of these gene promoters were occupied by H3K4me2 in the four-cell and eight-cell stages and the ICM; these genes played key roles in RNA metabolism and RNA splicing (Figure 7E). We then combined previous H3K4me3 data and chromosome accessibility data and found that these regions overlapped with high CG density regions, H3K4me3 enrichment, and chromatin accessibility regions (Figure 7E). Our results showed that H3K4me2 preferentially occupied hypomethylated and CpG-dense regions (Figures S5A-C).

**Figure 7.**
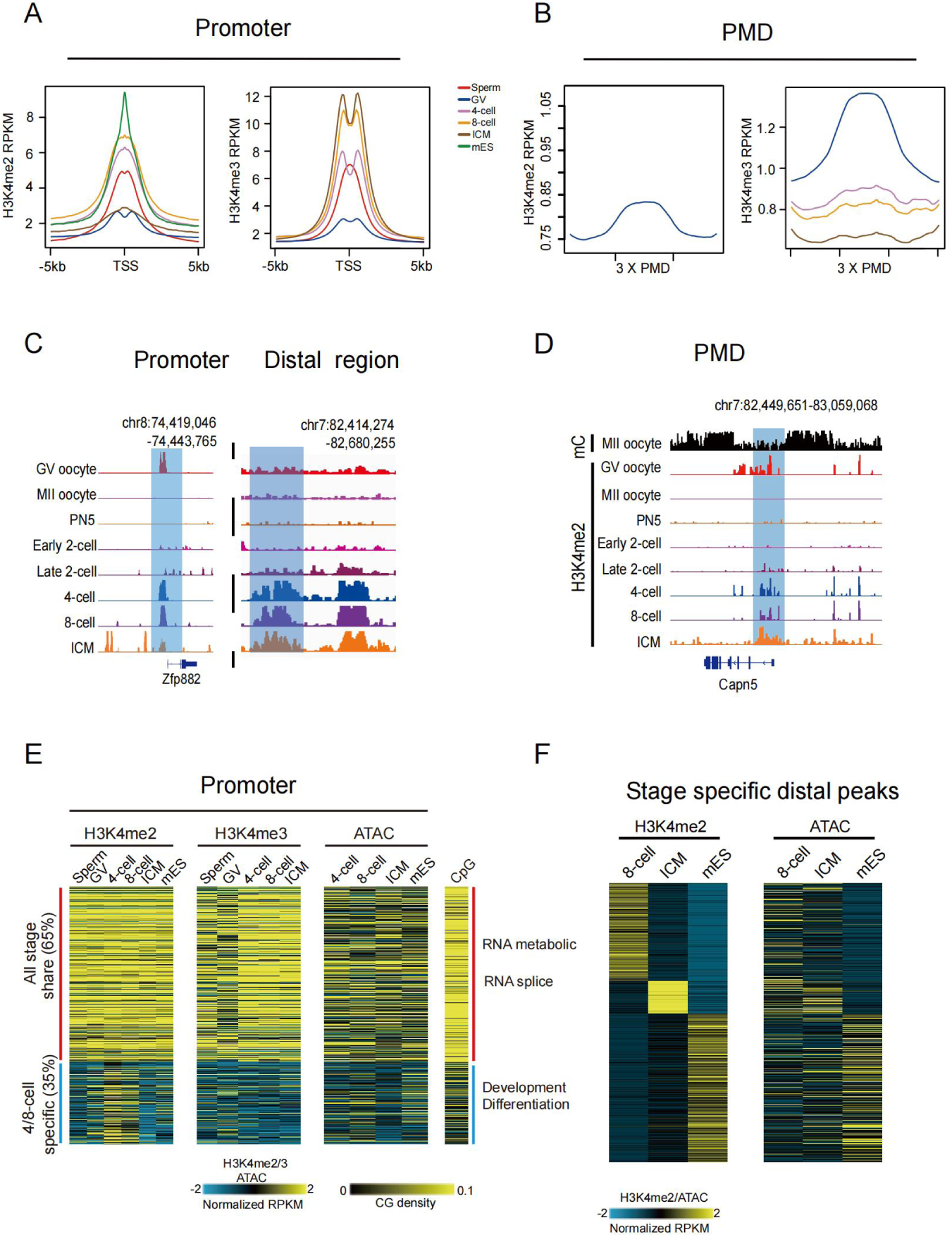
Pervasive H3K4me2 in CpG-rich regulatory regions in mouse embryos. (A-B) H3K4me2 and H3K4me3 enrichment near TSS (TSS ±5 kb) and PMD (3× PMD) in mouse gametes, pre-implantation embryos, and mESC. (C) The IGV view shows promoter H3K4me2 signals and widespread distal H3K4me2 signals in mouse oocytes and early embryos. (D) The IGV view shows H3K4me2 signals and the distribution of DNA methylation in mouse oocytes and early embryos at the PMD (partially methylation district) region. (E) Heat maps show the enrichment of H3K4me2 and H3K4me3 at the promoters, including that in all stages shared and four-cell/eight-cell specific. ATAC-seq, promoter CpG density, and GO analysis results are presented. (F) Heat maps of the distal peaks of H3K4me2 expressed in specific stages among eight-cell, ICM, and mESC. ATAC-seq are shown.

We further examined how the stage-specific distal peaks were reprogrammed and found that about 35% of the promoters of the genes that were associated with cell differentiation and development did not undergo the large-scale re-established of H3K4me2 in the four-cell, eight-cell, and blastocyst stages (Figure 7E). We identified genes expressed in specific stages and examined their promoter H3K4me2 (Figure S5E). We found promoters with constantly high H3K4me2 that are preferentially CpG-rich, whereas promoters with constantly low H3K4me2 were generally CpG-poor (Figures 7F and S5E). A group of genes showed dynamic promoter H3K4me2 that correlated with gene expression (Figure S5E). These genes played key roles in development, differentiation, and morphogenesis. Therefore, our results indicated that H3K4me2 of the promoter during early embryonic development correlated with gene activities and CpG density.

Next, we investigated distal H3K4me2 signals and found that four/eight-cell embryos also showed widespread distal H3K4me2 signals (Figure 7C). We first identified the distal H3K4me2 regions in mouse gametes and pre-implantation embryos. These regions were relatively CpG-rich (although lower than promoters) and hypomethylated (Figures S5B and S5C). However, compared to promoters, distal H3K4me2 was correlated with CpG density in a continuous and monotonic manner (Figure S5A). To summarize, these results indicated that widespread distal H3K4me2 occurred in CpG-rich and hypomethylated regions in mouse four/eight-cell embryos.

### Transcription regulation, CcREs priming, and resolving during maternal-to-zygotic transition

A study showed that most of the transcription factor binding regions overlap with H3K4me2-enriched regions^56^. To investigate the association between H3K4me2 and transcription regulation, we identified stage-specific distal regions and the accessibility in these distal peaks of H3K4me2 (Figure 7F). Then, we identified the binding motifs for transcription factors (TFs) enriched in these distal regions (Figure S5D). TFs in mESC and eight-cell embryos include pluripotency factors involved in the development of the system^57,58^. We found that CTF was specifically enriched in eight-cell embryos. The GATA and KLF families are enriched in ICMs.

In a study on humans, we found widespread deposition of H3K4me3 at the promoters, including those for developmental genes, before ZGA. These genes were either activated or repressed after ZGA^50^. Such promoter priming is related to embryonic development. To determine whether this also occurs in mice, we identified specific H3K4me2 target genes in the four/eight-cell stage and compared them to genes that are expressed or inactive at all stages. The H3K4me2 of promoters in the four/eight-cell-specific group decreased in ICM (Figures 8A and S6A). The “active” genes showed strong H3K4me2, H3K4me3, and ATAC-seq signals before and after ZGA (Figures 8A and S6A). Next, we investigated if such priming-resolving epigenetic transition also occurs in the distal regions. Distal H3K4me2 sites were enriched on regulatory elements, as 75.4% sites (compared to 16.2% of random) of them overlapped with candidate cis-regulatory elements (CcREs), identified by ENCODE (Figure 8B).

**Figure 8.**
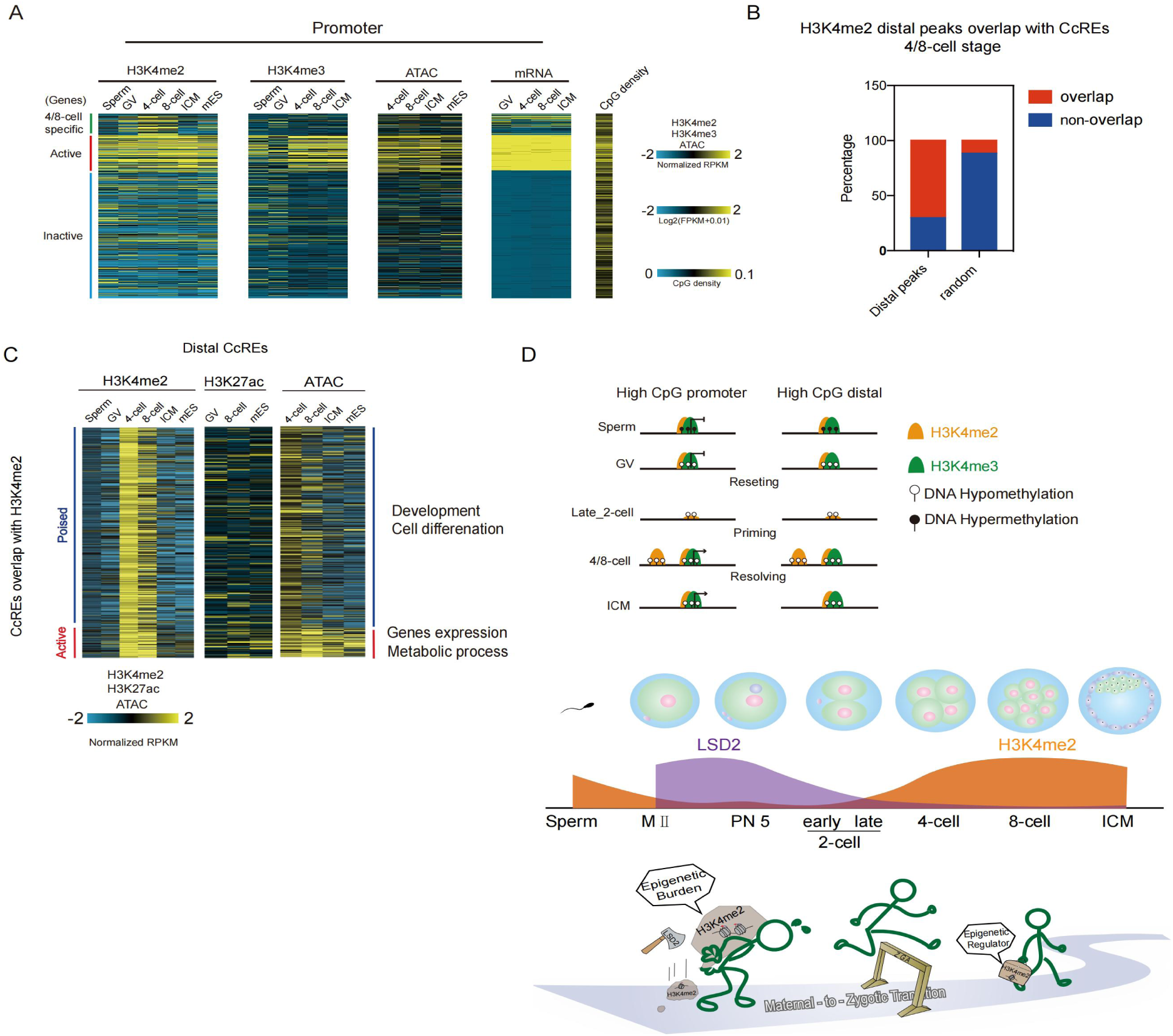
Chromatin states at specific stages and the H3K4me2 mode chart during mouse maternal-to-zygotic transition. (A) Heat maps of H3K4me2, H3K4me3, and ATAC-seq are shown, “All active” and “All inactive” promoters and the corresponding gene expression are presented. The CpG densities are also shown. (B) The bar graph shows that the H3K4me2 distal peaks overlapped with the CcREs four/eight-cell stage. Random peaks are shown as a control. (C) Heat maps of the chromatin states of the overlap between CcREs and the distal peaks of H3K4me2 and H3K27ac are presented. Clustering was conducted using ATAC-seq, H3K4me2 in the four-cell, eight-cell, and ICM stages. The results of the GREAT analysis for each cluster are shown (right). Data on histone modification of H3K27ac in mice was obtained from the GEO database, GSE72784. (D) Dynamic reprogramming of H3K4me2 during mouse parental-to-zygotic transition. A schematic model of “H3K4me2 resetting” during parental-to-zygotic transition is shown. After fertilization, the parental H3K4me2 is almost globally erased. *De novo* H3K4me2 promoters and distal regions in CpG-rich and DNA hypomethylated regions are re-established at the late two-cell stage, probably as a priming chromatin state. This leads to the resolution of primed promoters to “active” promoters with H3K4me3, or “poised”. A similar transition occurs for putative enhancers.

Enhancers are DNA sequences that increase promoter activity and, in turn, the frequency of gene transcription. The enhancer region is usually modified by H3K27ac (catalyzed by P300/CBP) and H3K4me1 histone 2 (catalyzed by MLL3/4)^59^, whereas the H3K4me3 modification level is low. This modification pattern is the opposite of the pattern found in the promoter region. We compared histone modification to H3K4me2 and H3K27ac and found no significant signal overlap between the two histone modifications at specific developmental stages in the distal region. Distal four/eight-cell H3K4me2 marked CcREs showed high enrichment of chromatin accessibility (Figures 8C and S6B). After the four/eight-cell stage, one class (“active”) remained accessible and was associated with basic biological processes (GREAT analysis). In contrast, the other class (Figures 8C and S6B; “poised”) was associated with H3K27me3 in the ICM and somatic cells; it was located preferentially near developmental genes. These results indicated that the widespread *de novo* H3K4me2 was re-established following ZGA, and H3K4me2 might be globally reset to a permissive state before it is resolved after the four/eight-cell stage (Figure 8D).

## DISCUSSION

Histone modifications are fundamental epigenetic marks. The reprogramming of histone marks plays an important role in early development^9^. In this study, we profiled the H3K4me2 landscape in mESC, oocytes, and early embryos using a modified low-input, high-sensitivity CUT&RUN method. We found the H3K4me2 showed differential patterns in sperms and oocytes and exhibited non-canonical patterns in GV oocytes but canonical patterns in the sperm in mice^36^. Additionally, the GV H3K4me2 peaks were almost erased in MII oocytes, and reconstructed accompanying ZGA occurred. Our results indicated that maternal H3K4me2 was lost during oocyte maturation. The non-canonical H3K4me2 in mammalian embryos and pervasive H3K4me2 in CpG-rich regulatory regions were found in four-cell and eight-cell embryos and in ICM. Hence, we described the first landscape and elucidated the potential function of H3K4me2 during early mammalian development. We also proposed a novel model, known as “Epigenetic Burden”, according to which parental H3K4me2 exhibits an “Epigenetic Burden” of maternal-zygote-transition. This burden is erased by maternal factor LSD2 before ZGA. H3K4me2 plays a regulatory role after ZGA (Figure 8 X). H3K4me2 undergoes a transition that starts from an “Epigenetic Burden”, and then becomes an “Epigenetic regulator” during early histone modification reprogramming.

The paternal epigenetic inheritance is important for development and disease in offspring. Significant epigenome reprogramming occurs and is associated with key processes during early mammalian development^1,3,9,14,39-41,60-64^. Genome-wide H3K4me3 exhibited a non-canonical pattern (ncH3K4me3) in full-grown and mature oocytes, i.e., it occurred as broad peaks at promoters and many distal loci^40,44,48^. We found that H3K4me2 was a non-canonical pattern in the GV oocyte, but it was weakened in the MII oocyte, indicating that H3K4me2 plays an important role during oocyte maturation. H3K4me3 is inherited in pre-implantation embryos and is erased in the late two-cell embryos when the process of canonical H3K4me3 is initiated^40,44^. The re-establishment of H3K4me2 mainly starts in the late two-cell stage, and it exists as a non-canonical pattern in the following stages. Several studies have shown that in many species, including mice, zebrafish, and humans, the parental epigenome undergoes dramatic reprogramming after terminally differentiated gamete fertilization, followed by subsequent re-establishment of the embryo epigenome, which leads to epigenetic ‘‘rebooting’’^9^. We found that H3K4me2 was erased by LSD2 after fertilization and re-established at the late two-cell stage. Inhibition of the deletion of H3K4me2 resulted in the arrest of embryonic development and abnormal expression of genes related to ZGA. This indicated that H3K4me2 is involved in the reactivation of the embryonic epigenome, and parental H3K4me2 is an “Epigenetic Burden”. A previous study reported strong H3K4me2 staining in zygotes and two-cell embryos using different immunofluorescence protocols^51^. This apparent discrepancy likely reflects differences in antibody sensitivity, confocal acquisition parameters, or embryo fixation conditions. In our hands, when we specifically adjusted the laser power and gain for the zygote and two-cell stages, weak H3K4me2 signals could be detected (data not shown). However, to maintain consistent imaging conditions across all stages from GV to blastocyst, we used identical parameters, which may have underestimated the signal at early stages. Therefore, our conclusion that H3K4me2 is largely erased and then re-established remains valid, but the precise dynamics may be more nuanced and dependent on technical settings.

H3K4me3 is an active gene expression marker, which mainly marks the gene promoter region. H3K4me1 and H3K4me2 mark enhancer and promoter regions. They have multiple functions for gene activation, according to different enzymes and complexes^22,30,35,65-74^. Some studies have reported a distinct pattern of H3K4me2 enrichment, spanning from the transcriptional start site (TSS) to the gene body, which marks a subset of tissue-specific genes. This pattern is associated with unique H3K4me2 profile mark tissue-specific gene regulation^33,35^. Our results showed non-canonical H3K4me2 in mammalian embryos and pervasive H3K4me2 in CpG-rich regulatory regions of four-cell and eight-cell embryos of mice. We next investigated whether H3K4me2 is associated with active or repressed genes in embryo development and found that promoter H3K4me2 in early mouse embryo development is correlated with gene activities and high CpG density. We also found that promoter H3K4me2 was consistent with H3K4me3.

Histone modification and chromatin accessibility profiles are usually used to identify transcription factor binding regions and motifs. In another study, we found large-scale chromatin accessibility in pre-ZGA human embryos, and these regions contained many transcription factors^3^. Around 90% of the transcription factor binding regions overlap with the H3K4me2-enriched regions. Stronger signals of H3K4me2 enrichment indicate a greater chance of overlap with the transcription factor binding regions^34^. In this study, we identified the binding motifs for transcription factors enriched in these distal regions, and the CCAAT transcription factor (CTF) was found to be specifically enriched in eight-cell embryos. The GATA and KLF families were enriched in ICMs. GATA is a key regulator of primitive endoderm (PE), which is a part of the ICM^75^, and KLF factors support naive pluripotency^76^. These results showed that stage-specific distal H3K4me2 involves transcription circuitry during mammalian embryonic development.

The state of chromatin influences DNA replication, DNA repair, and gene expression. Accessible chromatin occupies regulatory sequences, such as promoters and enhancers^77^. These elements interact with cell type-specific transcription factors to start transcription processes that determine cell fate^78^. Animal embryos in the early stages undergo extensive histone modification reprogramming, and chromatin remodeling before terminally differentiated gametes develop into pluripotent cells. Some studies have found extensive overlap of accessible promoters and enhancers at the four-cell stage in humans^50^. This finding indicated that crosstalk occurs between histone modification and the chromatin state. In our study, most of the H3K4me2 was re-established in chromatin-accessible regions, including promoters and distal regions at the four-cell stage. Thus, we inferred that H3K4me2 might play an important role in shaping the chromatin state. Our results indicated that crosstalk occurs between accessible chromatin and histone modification.

Dynamic histone modification plays an important role during mammalian maternal-to-zygotic transition (MZT). Approximately 22% of the oocyte genome associated with broad H3K4me3 is negatively correlated with DNA methylation and becomes confined to transcriptional-start-site (TSS) regions in two-cell embryos, accompanied by the onset of major ZGA^48^. Also, H3K4me2 is erased in MII oocytes, and sperm H3K4me2 is erased after fertilization. Similar to the pattern found for H3K4me3, H3K4me2 of the embryo was also re-established accompanied by the onset of major ZGA. The ncH3K4me3 in oocytes overlaps almost exclusively with partially methylated DNA domains^40^. H3K4me2 is also enriched in canonical gene promoters and partially methylated domains (PMDs).

A limitation of the current study is that we did not distinguish between maternal and paternal alleles when analyzing H3K4me2 distribution in pre-implantation embryos. Allele-specific H3K4me2 patterns could reveal asymmetric epigenetic inheritance and potential parental-origin effects on ZGA. Future studies using single - cell or low - input allele - resolved CUT&RUN, combined with SNP - based allele assignment, would be valuable to determine whether certain regulatory regions retain parental - specific H3K4me2 marks and how they contribute to embryonic development.

Overall, our findings provided a genome-wide landscape of H3K4me2 modification in mammalian oocytes and pre-implantation embryos. We showed that maternal LSD2 specifically demethylates H3K4me2 after fertilization in mammals. Also, H3K4me2 is erased to a greater extent in the male pronucleus than in the female pronucleus. Based on our results, we proposed a novel model, the “Epigenetic Burden” of histone modification reprogramming during mammalian development. This model can be used to further investigate the epigenetic regulation network in the early developmental stages of mammals.

## MATERIALS AND METHODS

### Culture of K562 cells

The human K562 cell line was cultured at 37 °C, 5% CO_2_, and saturated humidity. The cells were cultured in 1640 culture medium (GIBCO) containing 10% fetal bovine serum and 90% 1640 culture liquid.

### Cell culture of mouse ESC

Mouse embryo stem cells (ESC) were cultured on 0.1% gelatin-coated plates without irradiated mouse embryonic fibroblasts (MEFs), using standard ESC medium consisting of DMEM (GIBCO) supplemented with 15–20% fetal bovine serum (FBS, Sigma), 0.1 mM β-mercaptoethanol (Sigma), 0.1 mM MEM non-essential amino acids (Cellgro), 2 mM Glutamax (GIBCO), 1% nucleoside (Millipore), 1% EmbryoMax® Penicillin-Streptomycin Solution (Millipore), and 10 µg/L recombinant leukemia inhibitory factor (LIF) (Millipore). The mouse embryo stem cells were cultured at 37 °C, 5% CO_2_, and saturated humidity.

### Collection of mouse gametes and early embryos

The mouse oocytes were collected from 4–5-week-old C57BL/6N female mice (purchased from Charles River). The pre-implantation embryos were harvested from 4–5-week-old C57BL/6N females mated with DBA/2 males (eight-week-old when purchased from Charles River). The Germinal vesicle (GV) oocytes were obtained from 4–5-week-old C57BL/6N females, 46–48 h after they were injected with 10 IU of pregnant mare serum gonadotropin (PMSG, Solarbio). The 4–5-week-old C57BL/6N female mice were also administered 10 IU of human chorionic gonadotrophin (HCG) intraperitoneally, 46–48 h after PMSG was administered. Then, they were mated with DBA/2 males. The PN5 zygotes were collected at 28–32 h post-hCG injection. Early two-cell, late two-cell, four-cell, and eight-cell embryos, morulas, and blastocysts were harvested at 32–36 h, 45–48 h, 54–58 h, 60–64 h, 68–75 h, and 84–96 h post-hCG injection. The oocytes in the metaphase of second meiosis (MII) were collected 12–14 h after 10 IU of hCG was injected intraperitoneally. The zona pellucida of all oocytes and embryos was removed using glutathione (GSH Solarbio).

### Tranylcypromine treatment of mouse embryos

One-cell embryos were obtained at 24–26 h (PN3) and 28–32 h (PN5) post hCG injection and cultured in the G1 medium in the presence or absence of Tranylcypromine (5 mM). After 48 h, the embryos were transferred to the G2 medium with or without 5 mM TCP. All embryos were maintained at 37 °C in a humidified atmosphere containing 5% CO_2_. Then, morphologically normal PN5, late two-cell, and four-cell embryos were collected for further experiments.

### Immunofluorescence

The samples (mouse oocytes or pre-implantation embryos) were washed twice in MII medium (Sigma), and 4% PFA was used as a fixation buffer (BOSTER, AR1068) to fix samples at room temperature for about 60 min. Next, the samples were transferred using a mouth pipette and washed once with 1% BSA (Sigma, A1933) in 1×PBS (Corning, 21–040-CVR). The samples were then permeabilized using 0.5% Triton X-100 (Sigma, T8787) in 1×PBS at room temperature for 20 min. Then, they were washed with 1% BSA in 1×PBS and blocked using 1% BSA in 1×PBS at room temperature for 1 h. The blocked samples were incubated with H3K4me2 antibody (Sigma Millpore, 07030) in 1% BSA in 1×PBS diluted at a ratio of 1:500, at room temperature for 2 h or 4 °C overnight. After incubation, they were washed at least five times with 1% BSA in 1×PBS for 3 min. The samples were incubated with Alexa Fluor 568-conjugated donkey anti-rabbit IgG (Invitrogen, A10042) in 1% BSA in 1×PBS diluted at a ratio of 1:200, at room temperature for 50 min in the dark. After incubation, the samples were washed five times with 1% BSA in 1×PBS for 3 min. Finally, the samples were stained using the ProLong^@^ Gold Antifade Reagent with DAPI (Cell Signaling, 8961S), using a 4-Chamber Glass Bottom Dish (100 per case), 35 mm Dish with 20 mm Bottom Well (Cellvis, D35C4–20–1-N). The samples were incubated at room temperature for about 24 h in the dark. All images were captured using a Zeiss 700 confocal microscope. Raw images were projected along the Z-axis and processed using the ZEN software (Zeiss) with brightness and contrast adjusted (the same parameters were used for each channel of all samples). In the functional test section, we used the LSD2 antibody (Abcam, ab193080), H3K4me1 antibody (Abcam, ab176877), H3K4me2 antibody (mAb) (Active motif, 39079), and H3K4me3 antibody (Active motif, 39016) as primary antibodies (diluted at a ratio of 1:500). Secondary antibodies (diluted at a ratio of 1:300) used included Goat Anti-Rabbit lgG H&L (Alexa Fluor@488) (Abcam, ab150077) and Goat Anti-Mouse IgG H&L (Alexa Fluor® 488) (Abcam, ab150113).

### Zona pellucida removal and ICM/TE isolation

Before the CUT&RUN experiment was conducted, to guarantee the effect of permeation, the zona pellucida of all oocytes and embryos was chemically removed by glutathione (GSH Solarbio). The zona pellucida was removed after about 10–30 s. Then, the samples were washed twice in PBS (Corning 21–040-CVR) and washed once with wash buffer. The inner cell mass (ICM)s or TEs of day 4–5 blastocysts were isolated mechanically. Before performing CUT&RUN, the separated cells were washed twice with PBS and once with wash buffer, and then, resuspended in wash buffer.

### RNA-seq

For RNA-seq, 30–40 two-cell embryos were used for one sample. They were transferred to 1× PBS drops at room temperature and washed thrice to avoid any contamination. The cells were lysed in 1 µL of ice-cold lysis buffer for 10 min. Transcriptome amplification was performed using a Single Cell Full Length mRNA-Amplification Kit (Vazyme, N712). The RNA-seq library was constructed using True Prep DNA Library Prep Kit V2 for Illumina® (Vazyme, TD502). Finally, the library was sequenced using an Illumina Nova 6000 sequencer (Illumina). The data were analyzed following standard RNA-seq procedures.

### Generation of miniATAC-seq library

The miniATAC-seq libraries were prepared as previously described^3^. Cells for miniATAC-seq were prepared in the same way as those prepared for RNA-seq.

### CUT&RUN library construction and next-generation sequencing

The CUT&RUN method was used following published protocols^37,38^ with modifications in antibody binding, adaptor ligation, and DNA purification steps. All samples (including mouse zygotes, embryos with zona pellucida removed, or ESCs) were transferred into 0.2 mL conventional, non-low-binding PCR tubes (Axygen). The samples were resuspended in 60 µL of wash buffer (20 mM HEPES-KOH, pH = 7.5; 150 mM NaCl; 0.5 mM Spermidine; Roche complete protease inhibitor). Concanavalin-coated magnetic beads (Polyscience, 86057) were gently washed thrice or four times and resuspended in binding buffer (20 mM HEPES-KOH, pH = 7.9; 10 mM KCl; 1 mM CaCl_2_; 1 mM MnCl_2_), and then, 10–15 µL of Concanavalin-coated magnetic beads were added to each sample. The samples were incubated at room temperature (about 25 °C) at 450 rpm for 20 min in a Thermomixer (Eppendorf). The samples were placed on a magnetic stand for removing the supernatant, and then, resuspended in 50 µL antibody buffer (wash buffer plus digitonin 1000:1, 2 mM EDTA, pH = 8.0) with H3K4me2 antibody (Sigma Millpore, 07030) diluted at ratio of 1:100. The samples were incubated at room temperature (23–26 °C) for 2 h at 450 rpm on Thermomixer, or incubated overnight (≥ 16 h) at 4 °C and 450 rpm in a Thermomixer.

Next, the samples were held at the magnetic stand and washed with 190 µL of dig wash buffer (wash buffer plus digitonin, freshly pre-heated, 0.005%–0.01%; tested for each batch) once, and then, washed with 100 µL of the buffer again. The samples were then resuspended in 50 µL of dig wash buffer with PA/G-MNase (to a final concentration of 700 ng/mL), and incubated at room temperature (23–26 °C) on Thermomixer for 1 h, or incubated at 4 °C on Thermomixer for 3 h at 450 rpm. The samples were washed with 190 µL of dig wash buffer twice and then washed again with 100 µL of the buffer at the magnetic stand. Next, 100 µL of dig wash buffer was added and incubated on ice for 3 min. Targeted cleavage was performed on ice by adding 2 µL of 100 mM CaCl_2_ for 30 min, and the reaction was stopped by adding 100 µL of 2×stop buffer (340 mM NaCl; 20 mM EDTA, pH = 8.0; 4 mM EGTA, pH = 8.0; 50 µg/mL RNase A; 100 µg/mL glycogen; 0.005%–0.01% preheated digitonin; tested for each batch), and then, the mixture was fully vortexed. The samples were incubated at 37 °C for 30 min for fragment release. The total samples or supernatants were digested by adding 10 ng of carrier RNA (Qiagen, 1068337), 1 µL of 20% SDS, and 2 µL of Proteinase K (Roche). The reaction was performed at 56 °C for 45 min and 72 °C for 20 min in a thermal cycler. DNA was purified using phenol-chloroform, followed by ethanol purification. Then, 200 µL of phenol-chloroform was added, mixed, and centrifuged at 20 °C and 16,000 *g*for 5 min. Next, 200 µL of chloroform was added to each sample. After mixing, the samples were centrifuged at 20 °C and 16,000 *g* for 5 min and transferred to Lo-binding 1.5 mL tubes. Each sample was mixed with 2 µL of glycogen, 20 µL of NaOAC, and 500 µL of 100% ethanol (precooled at –20 °C). The mixture was incubated at –20 °C overnight.

The samples were centrifuged at 16,000 *g* and 4 °C for 20 min, and then, the supernatant was removed. Next, 1 mL of anhydrous ethanol was added to the samples, which were then centrifuged at 4 °C and 16,000 *g*for 10 min. The supernatant was removed by opening the lid of the tubes and air-drying. After the liquid disappeared, purified DNA was used for Tru-Seq library construction using NEB Next Ultra II DNA Library Prep Kit for Illumina (NEB, E7645S). For the NEB kit, the DNA was resuspended in 25 µL of ddH_2_O, followed by end-repair/A-tailing with 3.5 µL of End Prep buffer and 1.5 µL of an enzyme mix. The reaction was conducted at 20 °C for 30 min and 65 °C for 30 min in a PCR machine. Then, 1.25 µL of diluted adaptors (NEB Multiplex Oligos for Illumina, E7335S) were added and thoroughly vortexed. Next, 15 µL of Ligation master mix and 0.5 µL of Ligation enhancer were added, and the mixture was incubated at 16–20 °C for 16–20 min, or at 4 °C overnight. After the ligation reaction was over, the samples were treated with 1.5 µL of USERTM enzyme. The reaction was performed at 37 °C for 15 min in a PCR machine. The samples were purified using 1×AMPure beads. The PCR was performed by adding 25 µL of 2×KAPA HiFi Hot Start Ready Mix (KAPA biosystems, KM2602) with 2.5 µL of primers of the NEB Oligos kit and 2.5 µL of Illumina universal primers. The conditions used were as follows: 98 °C for 45s, (98 °C for 15s and 60 °C for 10s) with 16–17 cycles, and 72 °C for 1 min. The final libraries were purified using 0.9×AMPure beads and used for next-generation sequencing.

### Data analysis

#### CUT&RUN data processing

The single-end CUT&RUN data were mapped to the mm9 genome using Bowtie2 (Version 2.2.9)^79^ with the parameters –t –q –N 1 –L 25. The paired-end CUT&RUN data were mapped based on the parameters: -t –q –N 1 –L 25 –X 1000 --nomixed --no-discordant. All unmapped reads, non-uniquely mapped reads, and PCR duplicates were removed. For downstream analysis, the read counts were normalized by calculating the number of reads per kilobase of bin per million reads sequenced (RPKM) for 100-bp bins of the genome. The RPKM values were further Z-score normalized for minimizing the batch effect.

#### ATAC-seq data processing

The ATAC-seq reads were mapped to the mm9 genome using Bowtie2 (version 2.2.2)^79^ with previously described parameters. All unmapped reads, non-uniquely mapped reads, and PCR duplicates were removed. For downstream analysis, the read counts were normalized by calculating the RPKM for 100 bp-bins of the genome. The RPKM values were further normalized by Z-score normalization to minimize the batch effect.

#### DNA methylation data processing

We used the Bismark software for mapping DNA methylation reads to the mm9 genome. PCR duplicates were removed. For each CpG site, the methylation level was computed as the total methylated counts divided by the total counts across all reads covering the CpG site.

#### Analysis of the promoter H3K4me2 and H3K4me3 enrichment

Annotated promoters (TSS±2.5 kb) of the mm9 refFlat annotation database from the UCSC genome browser were used. Whole-genome Z-score normalized RPKM scores were used for assessing the data.

#### Identification of stage-specific distal ATAC-seq peaks

Stage-specific distal ATAC-seq peaks were identified as described previously. Briefly, distal peaks from human embryos and ES cells were merged, and then, the average Z-score was normalized. The RPKM scores were calculated based on them. A Shannon entropy-based method ^3^ was used to identify stage-specific peaks. Peaks with entropy less than 2 were selected as candidates and were further screened using the following criteria: the distal peak had high enrichment at this stage (normalized RPKM > 1) and positive enrichment (normalized RPKM > 0) at no more than two additional stages.

Minor/major ZGA genes and stage-specific genes were screened using the RNA-seq data from mouse germ cells and embryos. ZGA genes were defined as those unexpressed or underexpressed in oocytes (FPKM < 5) but highly upregulated (FPKM > 5, at least three-fold upregulation) in either the one-cell or early two-cell embryo stage (minor ZGA genes) or the late two-cell embryo stage (major ZGA genes). The genes that were highly expressed in oocytes and considerably upregulated (over five-fold upregulation) during the late two-cell embryo stage were also defined as major ZGA genes.

For stage-specific genes in eight-cell embryos and ICMs, a Shannon entropy-based method ^3^ was used to identify stage specific-genes, as previously described. Maternally expressed genes, defined as those expressed in GV or MII oocytes (FPKM ≥ 1), were excluded from this analysis to avoid the confounding effects of lingering maternal RNAs.

### Clustering analysis

We conducted k-means clustering of gene expression levels at various stages using Cluster 3.0 with the parameters –g 7 (Euclidean distance). Heat maps were generated using Java Treeview 3.0. We conducted hierarchical clustering analysis for H3K4me3 at various stages using the R package “ape” based on Pearson’s correlation between each pair of stages. The distance between the stages was calculated as 1 − Pearson’s correlation. Identification of oocyte PMDs We calculated the average DNA methylation levels for 10-kb bins of the genome and counted the number of CpG dyads in each 10-kb bin. Bins with an average DNA methylation level below 0.6 and above 20 CpG dyads were selected and merged into PMDs using BEDtools (Version v2.25.0)^80^. The promoter regions (±2.5 kb) were removed from PMDs.

### Identification of H3K4me2 peaks

The H3K4me2 peaks were called using MACS2 (Version 2.0.9)^81^ with the parameters --broad --nomodel --nolambda -g mm 1.87e9/-g hs 2.7e9. Noisy peaks with weak signals (summed RPKM < 0) were removed before further analyses were conducted.

### Statistical analysis

All statistical analyses in this study were briefly described in the text and performed using R (v3.6.1) (http://CRAN.R-project.org/).

## DATA AVAILABILITY

All raw sequencing data generated during this work have been submitted to the Genome Sequence Archive in National Genomics Data Center (China National Center for Bioinformation/Beijing Institute of Genomics, Chinese Academy of Sciences). The GSA accession ID of this paper is CRA004978.

## FUNDING AND ACKNOWLEDGEMENT

This work was supported by the National Key R&D Program of China (2019YFA0802200 and 2019YFA0110900 to Jiawei Xu), the National Natural Science Foundation of China 32170819 to Jiawei Xu, Henan University science and technology innovation team to Jiawei Xu), Scientific Research Innovation team of the First Affiliated Hospital of Zhengzhou University Excellent innovation team to Jiawei Xu, Henan Zhongyuan Medical Science and Technology Innovation Development Foundation to Jiawei Xu.

## Author contributions

Conceptualization: JWX,HC

Methodology: CW, YQW, YL

Experiments performing: CW, YL, YQW, KYH, XRM, WBL

Data NGS (next generation sequencing): CW,YL,YQW

Bioinformatics analysis: YS, JWX, CW, YL, JG Supervision: JWX

Writing—original draft: JWX, CW, YS, YQW, YL

Writing—review & editing: JWX

## Competing interests

The authors declare no competing interests.

## Extended Data

**Extended Data Figure 1.**
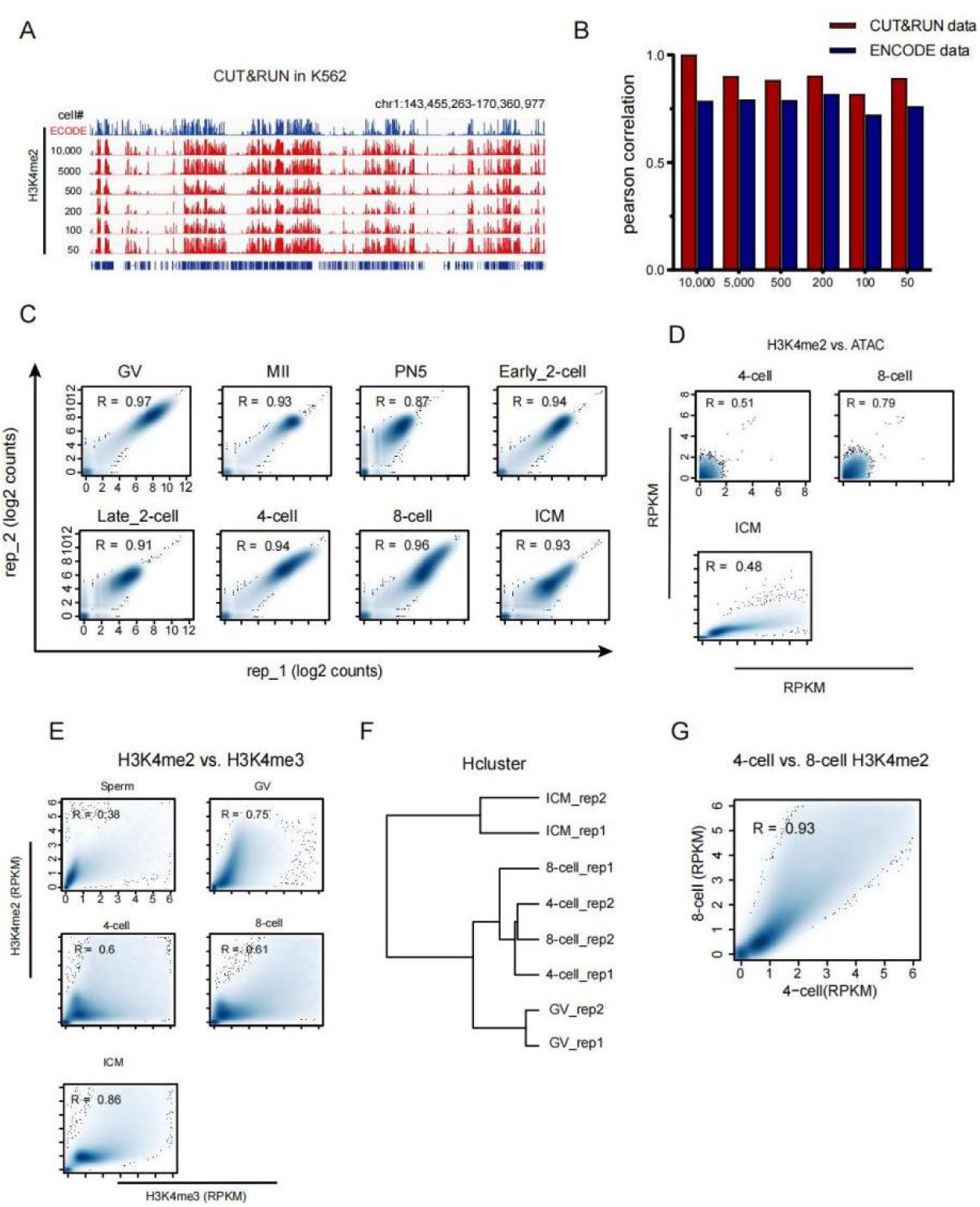
Validation of H3K4me2 CUT&RUN data in Human K562 cell line, oocytes and early embryos. (A) The IGV views showing H3K4me2 distributions by CUT&RUN using various numbers of Human K562 cell line. The ENCODE references are added for comparison. GEO accession: GSM733651. (B) The Pearson correlation coefficients showing the comparison between CUT&RUN of H3K4me2 using various numbers of Human K562 cells and conventional ChIP–seq data from ENCODE. (C) Scatter plots showing the correlations of biological replicates (n=2) of H3K4me2 signals (2kb-bin, whole genome) by CUT&RUN for each stage of mouse oocytes and early embryos. The Pearson correlation coefficients are also shown. (D) Scatter plots comparing the H3K4me2 signals with ATAC-seq (Assay for Transposase-Accessible Chromatin with high throughput sequencing) at promoters or distal accessible regions in mouse oocytes and early embryos. Spearman correlation coefficients are also shown. (E) Scatter plots comparing the H3K4me2 signals with H3K4me3 signals in mouse sperm, oocytes and early embryos. Spearman correlation coefficients are also shown. (F) Hierarchical clustering of H3K4me2 mouse oocytes and early embryos. Pearson correlation was used to measure distances. (G) Scatter plots comparing the H3K4me2 signals between 4-cell and 8-cell in mouse embryos. Spearman correlation coefficients are also shown.

**Extended Data Figure 2.**
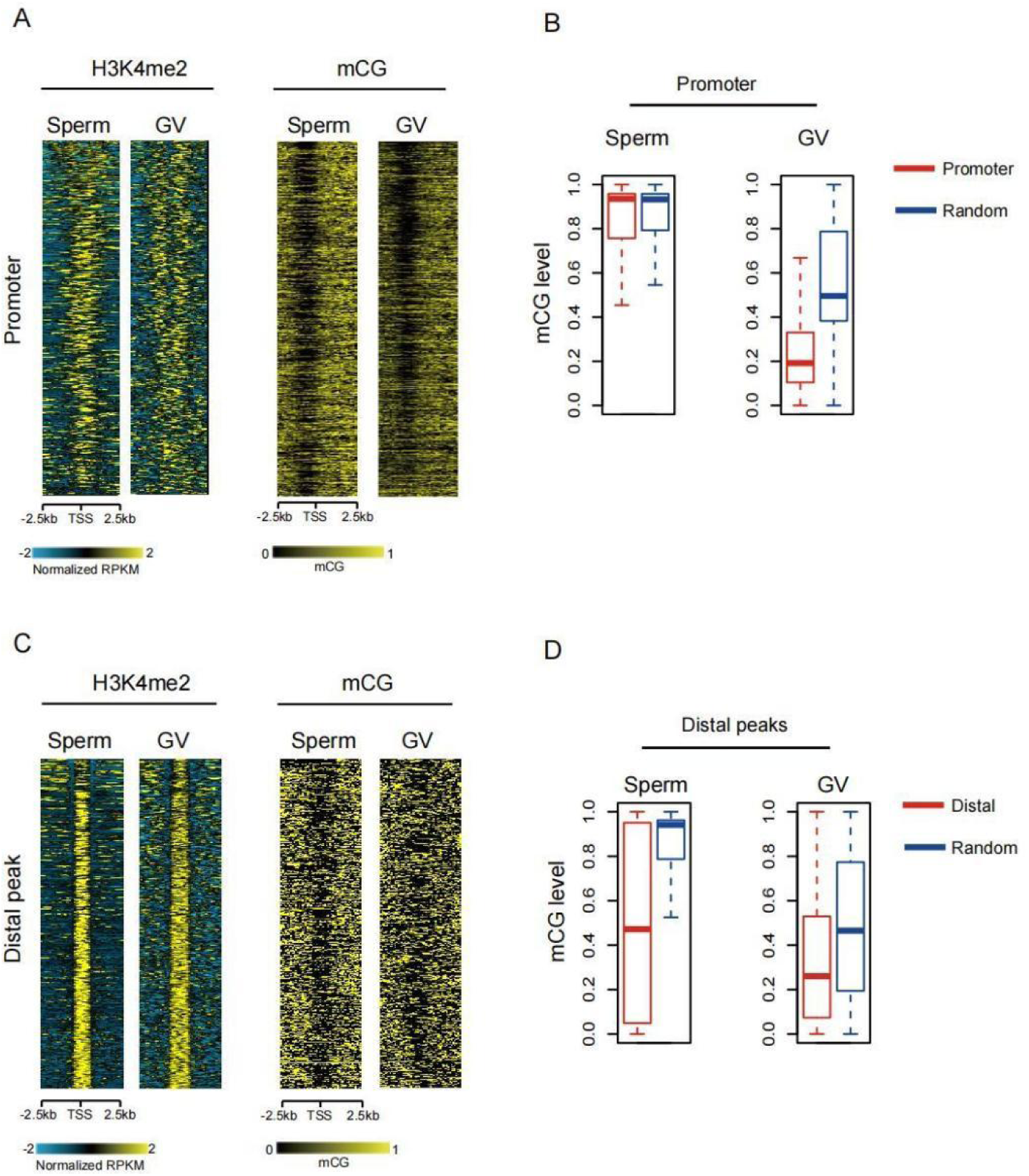
H3K4me2 in mouse gametes. (A) Heatmaps showing H3K4me2 signals at all promoter regions (TSS ± 2.5kb) in mouse gametes. DNA methylation (TSS ± 2.5kb) is also mapped. DNA methlation data cited in GEO dataset: GSE56697. (B) The box plots showing DNA methylation level at all promoter regions in mouse gametes. Random peaks are shown as a control. DNA methlation data cited in GEO dataset: GSE56697. (C) Heatmaps showing H3K4me2 signals at all distal regions (peak ± 2kb) in mouse gametes. DNA methylation (peak ± 2kb) is also mapped. DNA methlation data cited in GEO dataset: GSE56697. (D) The box plots showing DNA methylation level at all distal regions in mouse gametes. Random peaks are shown as a control. DNA methlation data cited in GEO dataset: GSE56697.

**Extended Data Figure 3.**
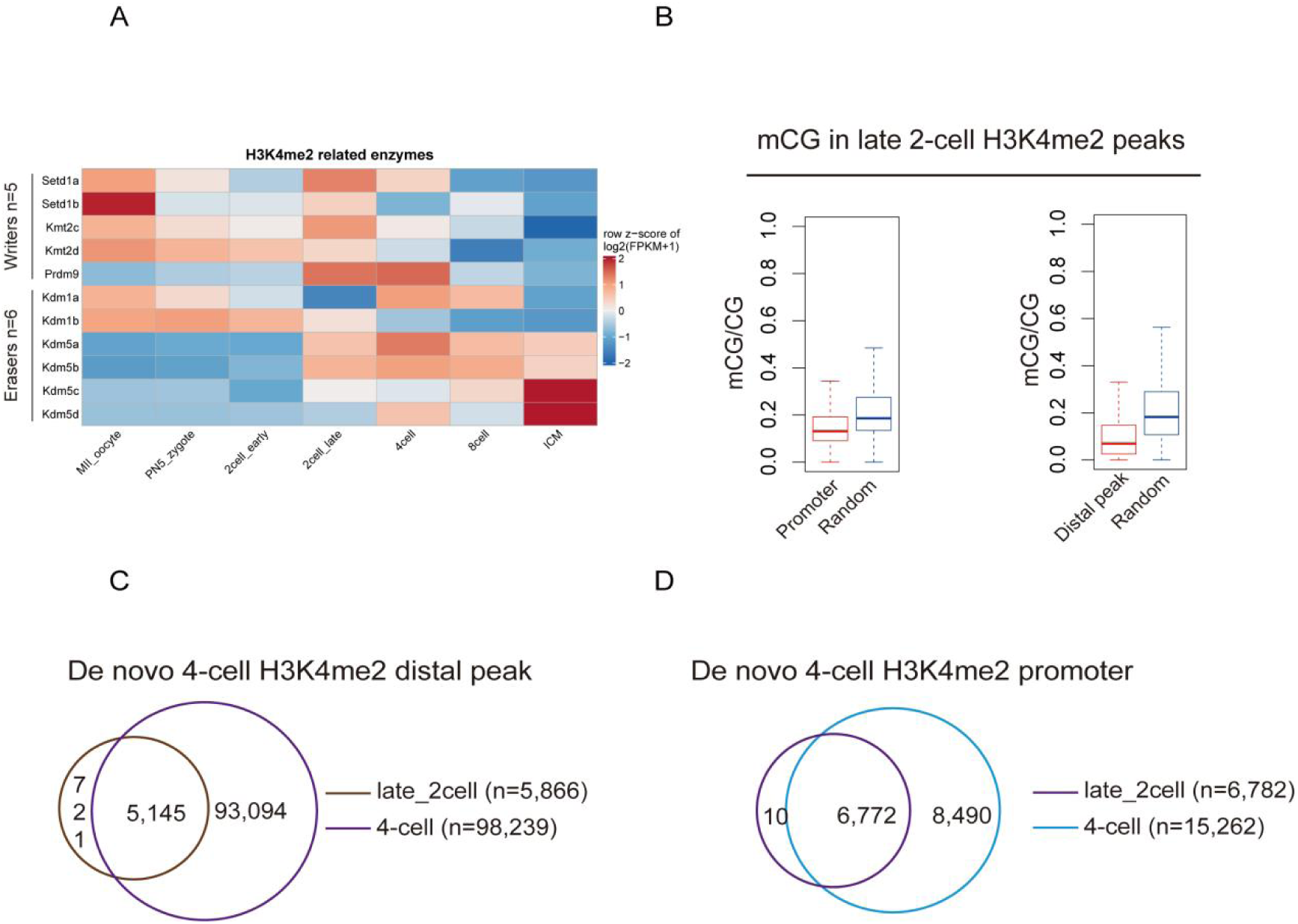
H3K4me2 in late 2-cell embryos. (A) Heatmaps showing the expression of H3K4me2 related regulators in mouse (based on published RNA-seq data43). (B) The box plots showing DNA methylation level at all promoter regions and distal region in mouse 2-cell stage. Random peaks are shown as a control. (C-D) Venn diagrams showing De novo 4-cell H3K4me2 promoter and distal peaks number.

**Extended Data Figure 4.**
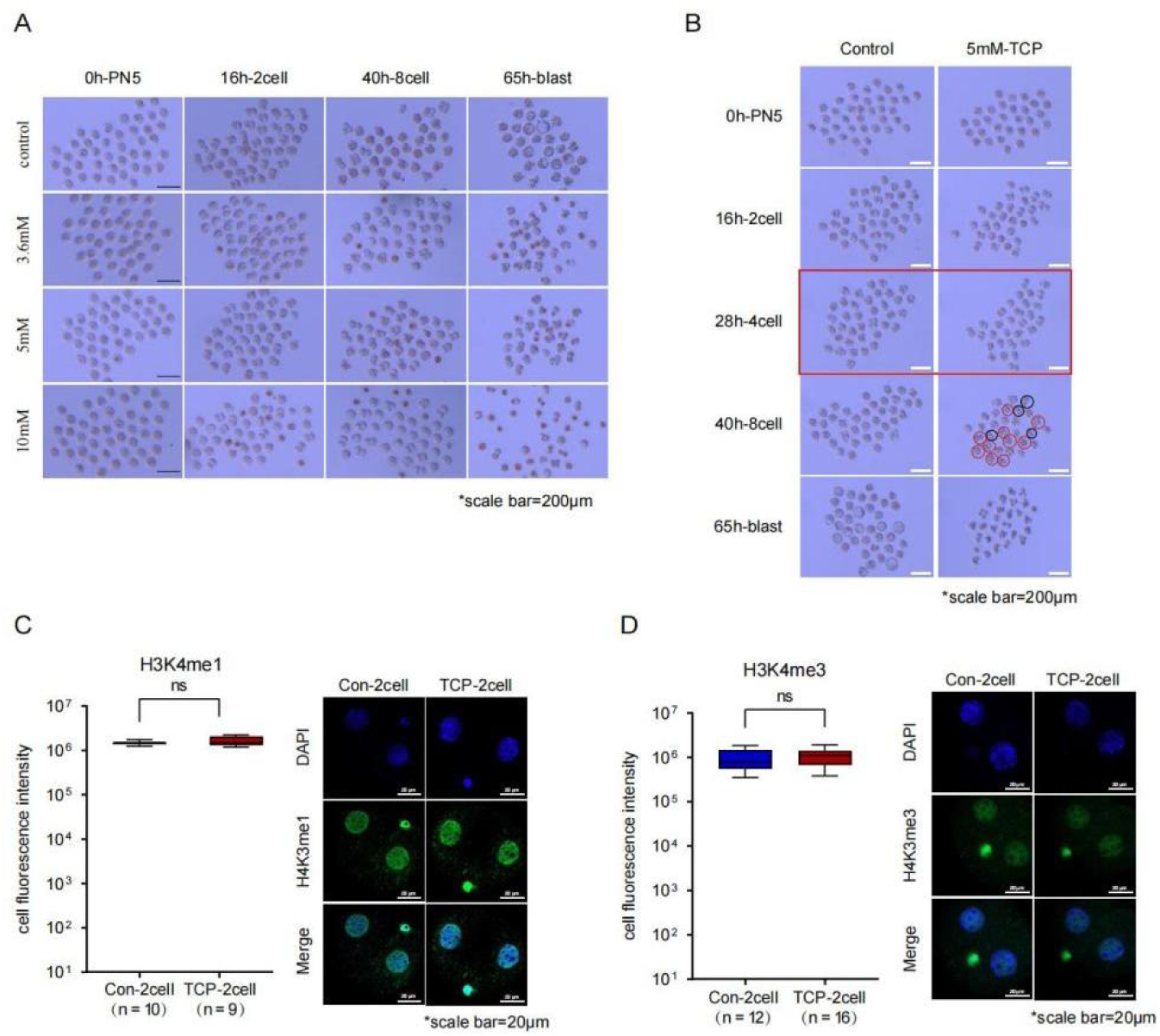
Cell culture experiment under TCP treatment. (A) Morphological observation of cells under different concentration of TCP culture, scale bar is also shown. (B) Morphological observation of embryo development after 5mM TCP inhibition of LSD2 expression. (C-D)Immunofluorescence images showed the immunofluorescence intensity of H3K4me1 (C) and H3K4me3 (D) of 2-cell embryos in the experimental group (5mM TCP) and control group, scale bar is also shown.

**Extended Data Figure 5.**
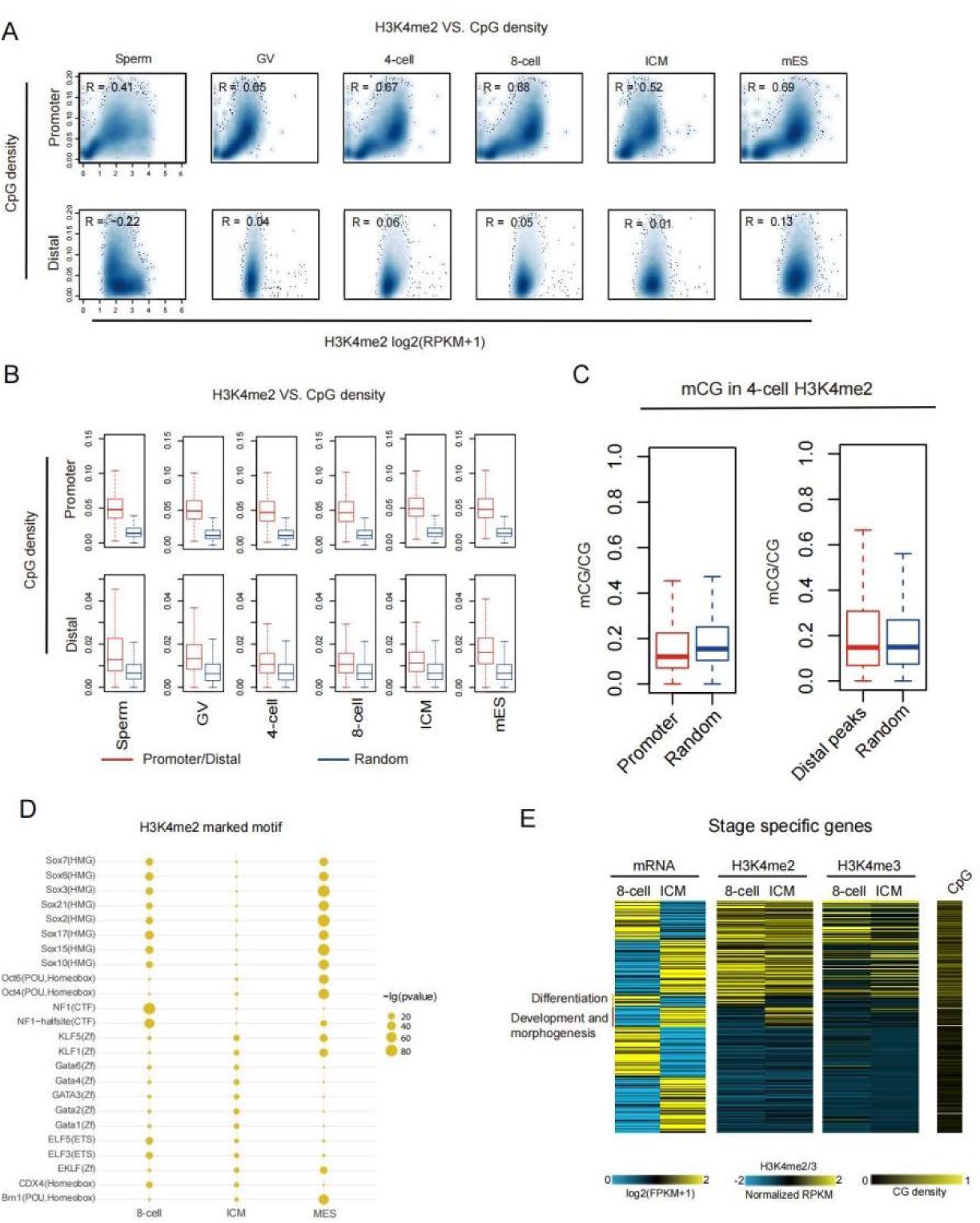
H3K4me2 in mouse 4/8-cell embryos. (A) Scatter plots comparing the promoter and distal H3K4me2 signals with CpG densities in mouse gametes, 4-cell, 8-cell, ICM and mESC. The Spearman correlation coefficients are also shown. (B) The box plots showing the CpG density and promoter/distal H3K4me2 peaks. Random peaks are shown as a control. (C) The box plot showing DNA methylation in promoter and distal 4-cell H3K4me2 peaks. Random peaks are shown as a control. (D) TF motifs identified from active enhancers in mouse early embryos (8-cell, ICM) and mESC. showing motif enrichment p value < 1e-20 at least at one stage were included. Circle size showing TF enrichment and the expression of the TF is color coded. (E) Heatmaps showing stage-specifically expressed genes further classified by their H3K4me2 states (high, dynamic, and low). RNA-seq, H3K4me3 state, CpG densities and GO analysis results are also mapped and shown.

**Extended Data Figure 6.**
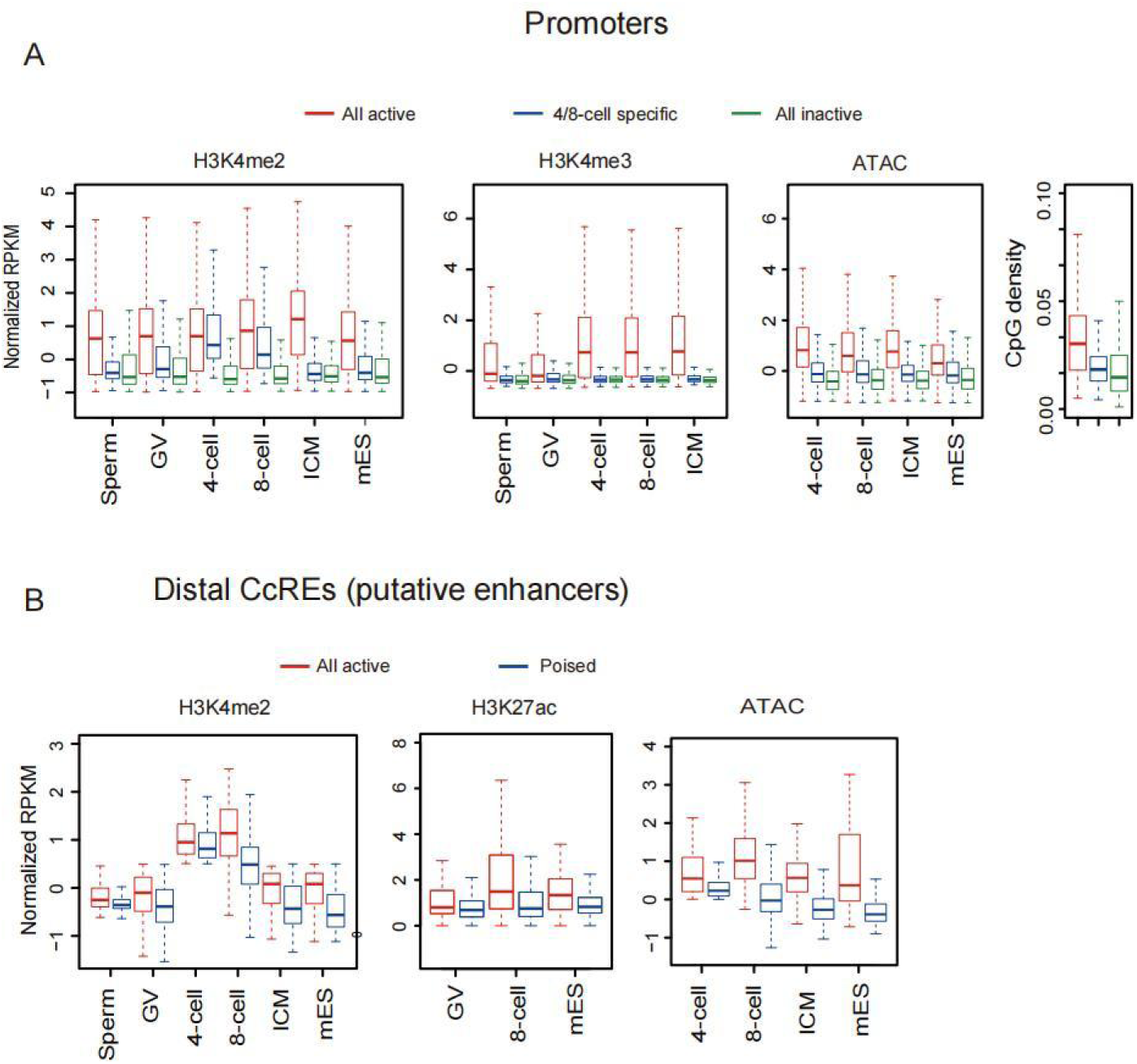
Chromatin states at specific stages and distal CcREs. (A) Box plots showing H3K4me2, H3K4me3 and ATAC-seq signals, “All active” and “All inactive” promoters in mouse gametes, embryos and mESC. CpG density for each group is also shown (right). (B) The box plot showing the ATAC-seq, H3K4me2 and H3K27ac signals at putative active, poised or random enhancers in mouse gametes, early embryos or mESC, Mouse histone modified H3K27ac data refer to GEO database: GSE72784.

## Extended Data Table

The sample information in this study was listed in the table below.

**Table 1.**
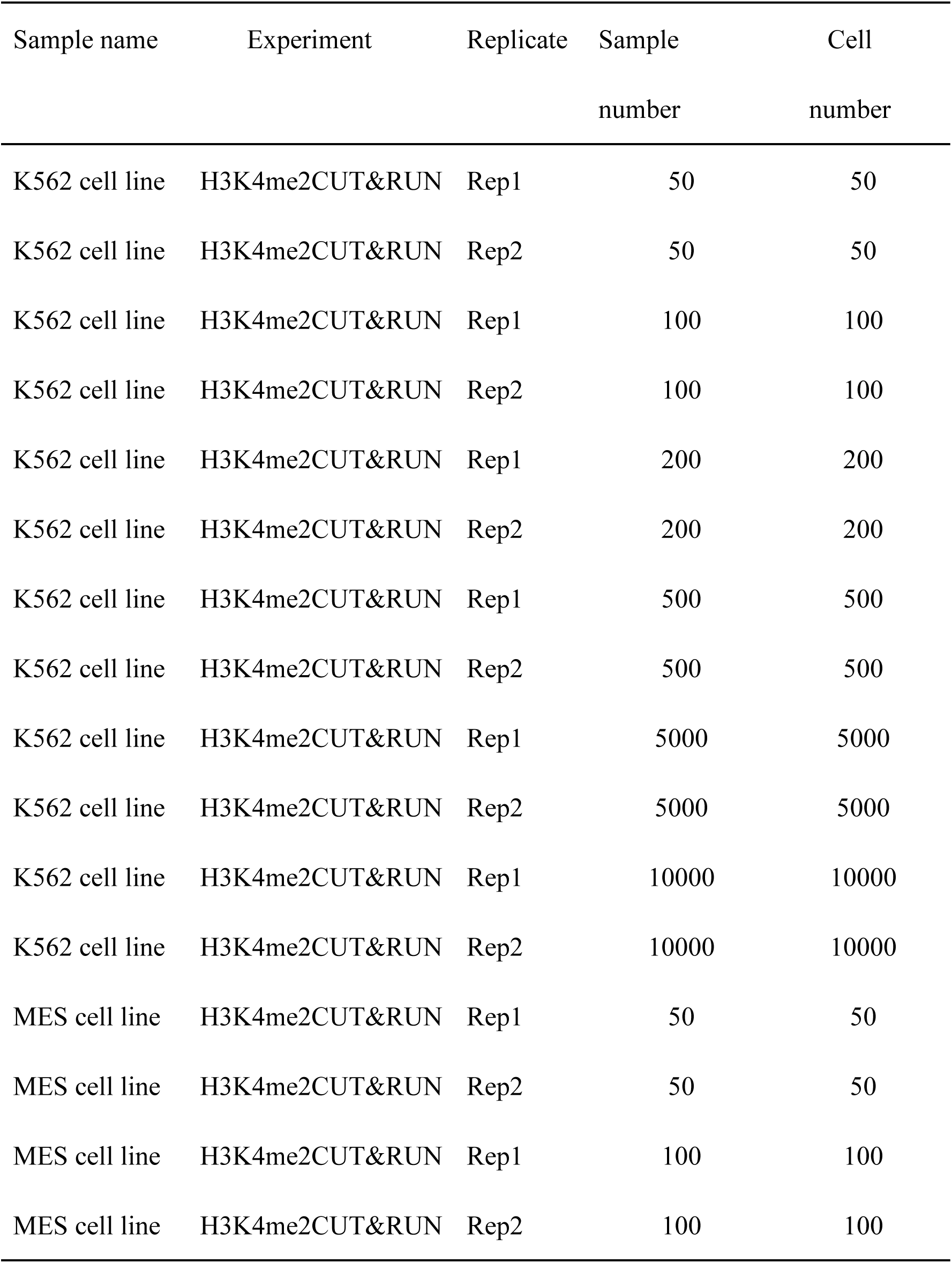
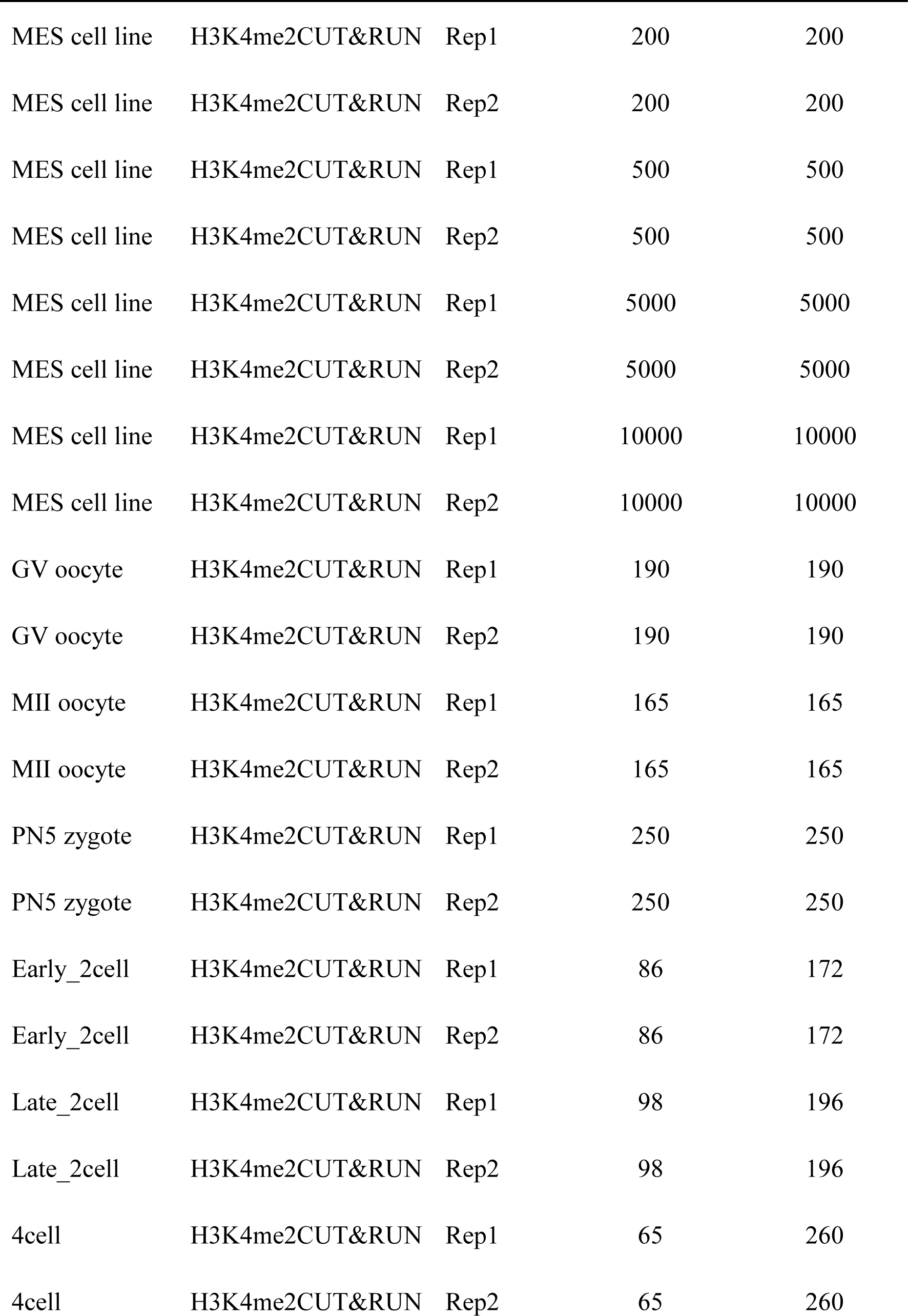
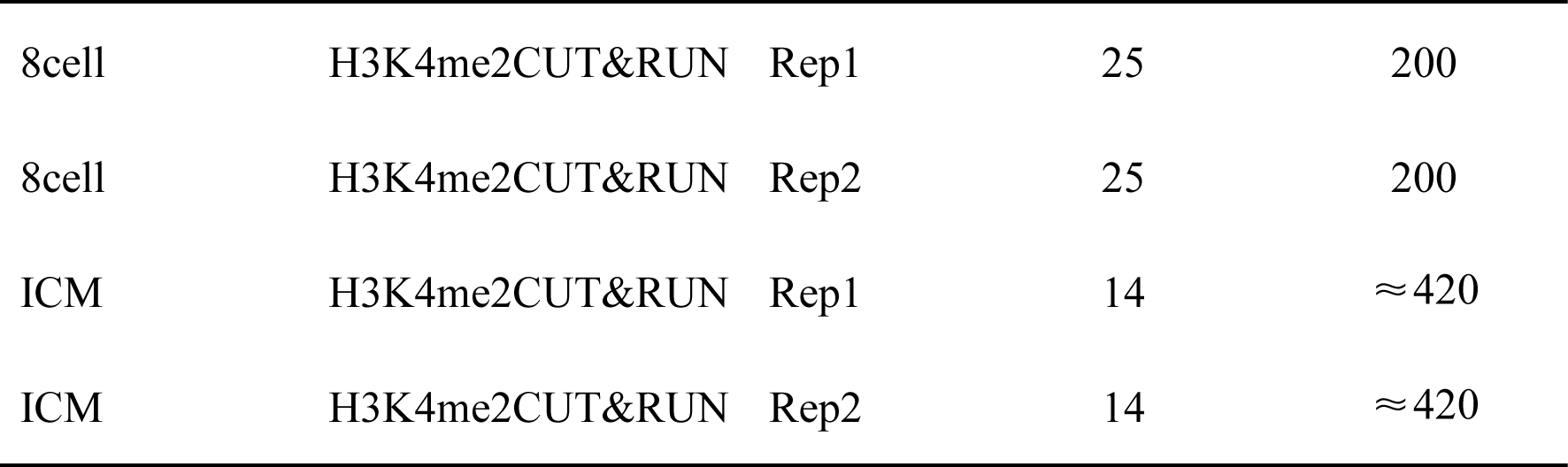
CUT&RUN experimental sample information.

## Notes

https://ngdc.cncb.ac.cn/gsub/submit/gsa/subCRA006991/finishedOverview

https://ngdc.cncb.ac.cn/gsa/

### Competing Interest Statement

The authors have declared no competing interest.

### Summary of Updates

In response to the reviewer comments, we have made the following revisions to the manuscript. Modification 1: (related to H3K4me2 staining discrepancy with Ancelin et al., 2016) was added in both the Results and Discussion sections, explaining that differences in confocal parameters, antibody sensitivity, or fixation conditions may affect signal detection, and that weak signals could be observed when acquisition settings were optimized for zygote/2‑cell stages. Modification 2：addressed missing references on page 9: we provided an empirical explanation for the TCP concentration (2a), and added existing citations [1,3,9] to the sentence on histone modification regulation (2b) as well as citation [55] to the Zfp825 statement (2c). Modification 3：added a new paragraph in the Discussion acknowledging the lack of allele‑specific H3K4me2 analysis and recommending future studies using allele‑resolved CUT&RUN. Modification 4: revised overstatements throughout the manuscript to reflect correlation rather than causality (e.g., alters → was associated with altered, can affect → could affect, inhibits → may inhibit). Modification 5：corrected the misinterpretation of LSD2 expression data: we now accurately state that LSD2 is highly expressed in growing oocytes but decreases in MII oocytes, with the heatmap re‑normalized. One new reference (Ancelin et al., 2016) was added to the reference list. All changes have been highlighted and labelled accordingly.

