## Supplemental figures for "Resetting of H3K4me2 during mammalian parental-to-zygote transition"

### Extended Data

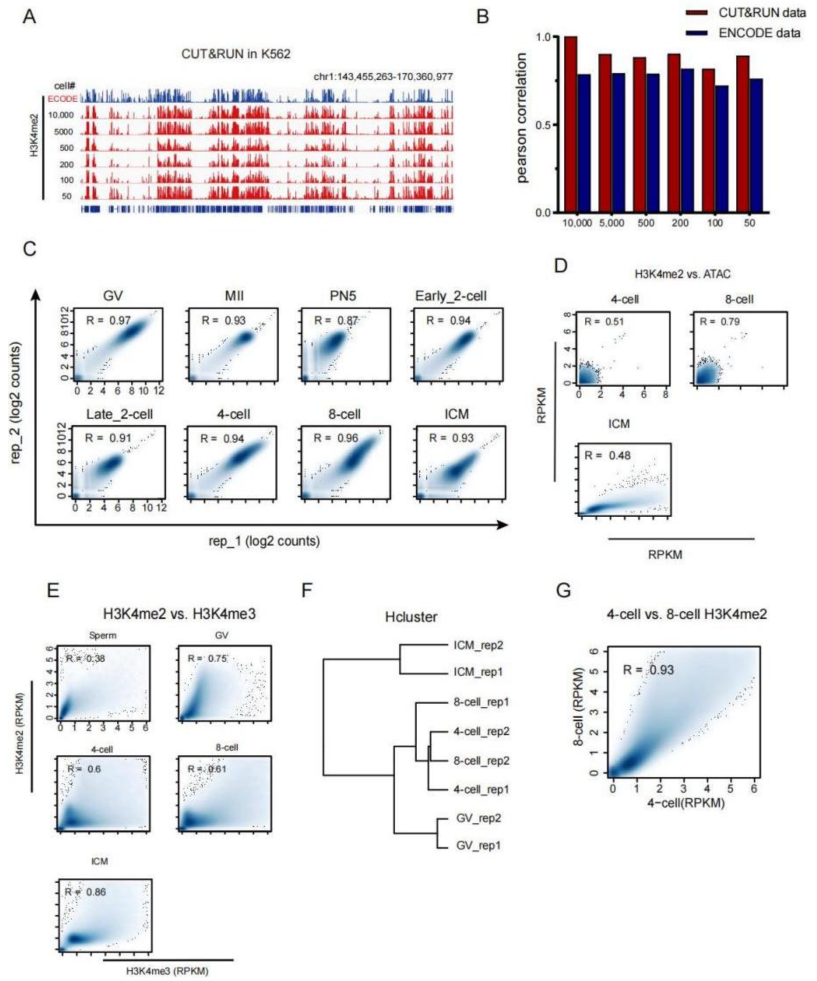


**Extended Data Figure 1. Validation of H3K4me2 CUT&RUN data in Human K562 cell line, oocytes and early embryos.**

(A) The IGV views showing H3K4me2 distributions by CUT&RUN using various numbers of Human K562 cell line. The ENCODE references are added for comparison. GEO accession: GSM733651 .

(B) The Pearson correlation coefficients showing the comparison between CUT&RUN of H3K4me2 using various numbers of Human K562 cells and conventional ChIP–seq data from ENCODE.

(C) Scatter plots showing the correlations of biological replicates (n=2) of H3K4me2 signals (2kb-bin, whole genome) by CUT&RUN for each stage of mouse oocytes and early embryos. The Pearson correlation coefficients are also shown.

(D) Scatter plots comparing the H3K4me2 signals with ATAC-seq (Assay for

Transposase-Accessible Chromatin with high throughput sequencing) at promoters or distal accessible regions in mouse oocytes and early embryos. Spearman correlation coefficients are also shown.

(E) Scatter plots comparing the H3K4me2 signals with H3K4me3 signals in mouse sperm, oocytes and early embryos. Spearman correlation coefficients are also shown.

(F) Hierarchical clustering of H3K4me2 mouse oocytes and early embryos. Pearson correlation was used to measure distances.

(G) Scatter plots comparing the H3K4me2 signals between 4-cell and 8-cell in mouse embryos. Spearman correlation coefficients are also shown.

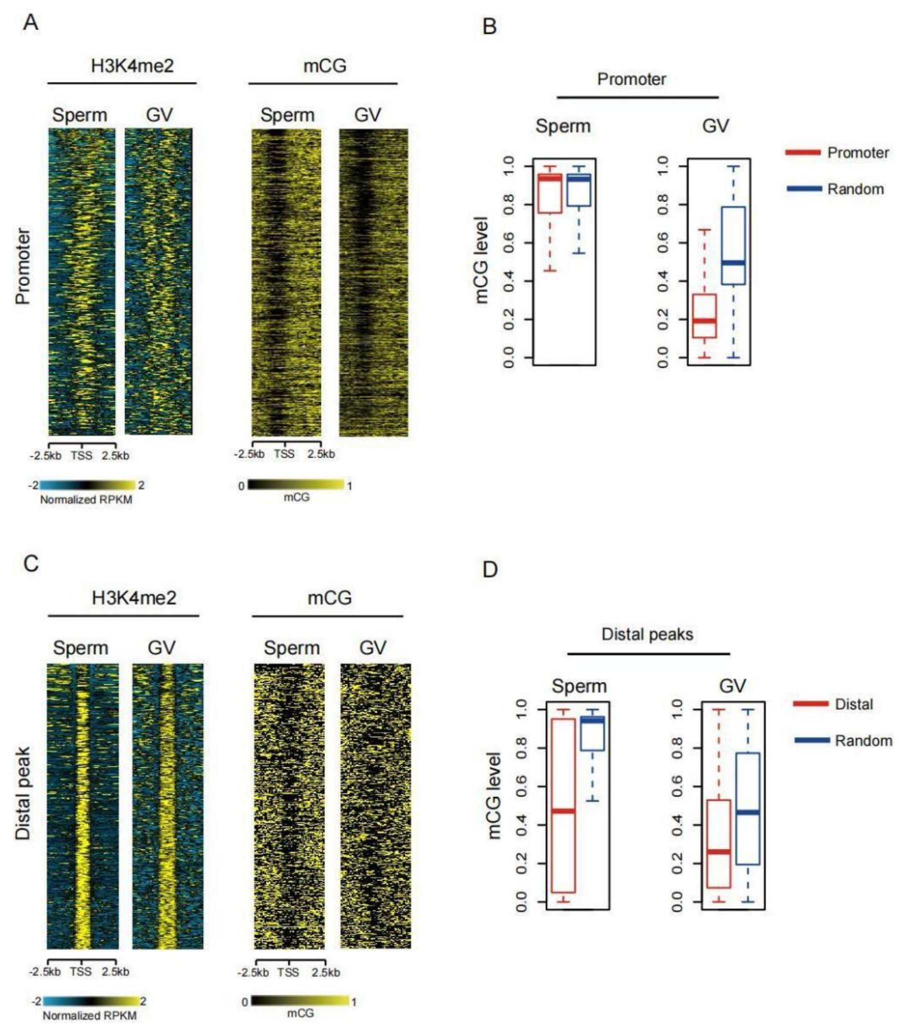


**Extended Data Figure 2. H3K4me2 in mouse gametes.**

(A) Heatmaps showing H3K4me2 signals at all promoter regions (TSS ± 2.5kb) in mouse gametes. DNA methylation (TSS ± 2.5kb) is also mapped. DNA methlation data cited in GEO dataset: GSE56697.

(B) The box plots showing DNA methylation level at all promoter regions in mouse gametes. Random peaks are shown as a control. DNA methlation data cited in GEO dataset: GSE56697.

(C) Heatmaps showing H3K4me2 signals at all distal regions (peak ± 2kb) in mouse gametes. DNA methylation (peak ± 2kb) is also mapped. DNA methlation data cited in GEO dataset: GSE56697.

(D) The box plots showing DNA methylation level at all distal regions in mouse gametes. Random peaks are shown as a control. DNA methlation data cited in GEO dataset: GSE56697.

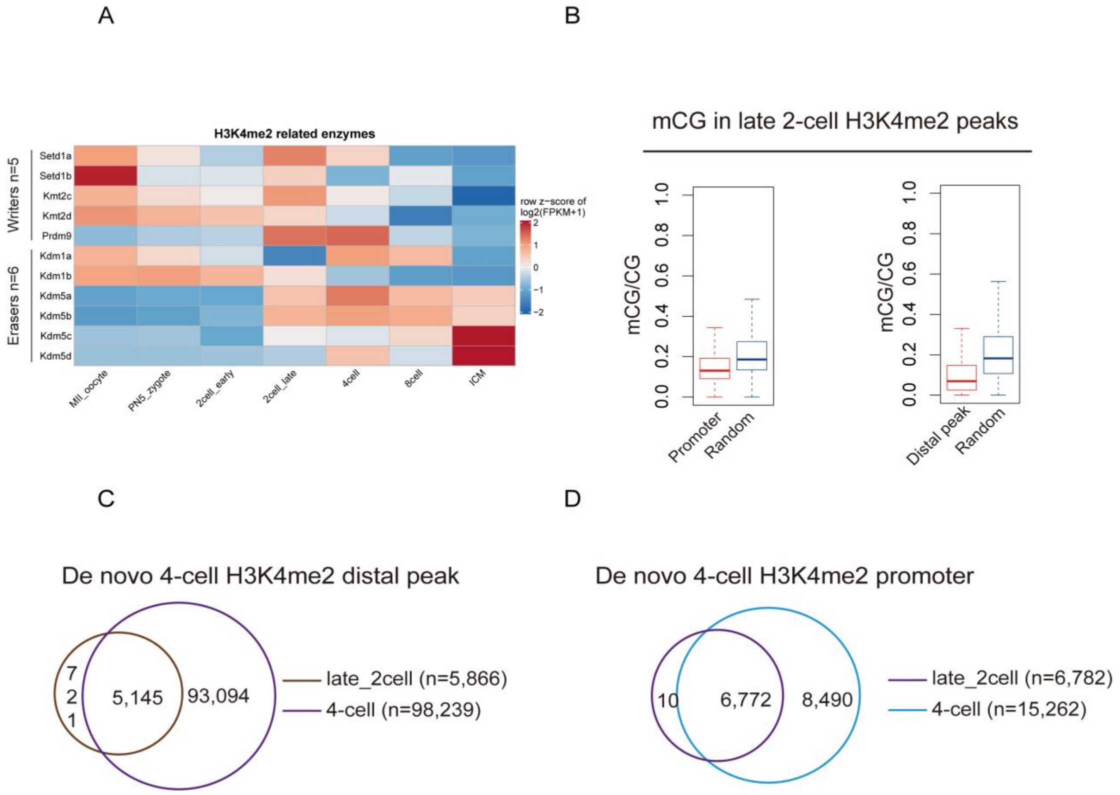


**Extended Data Figure 3. H3K4me2 in late 2-cell embryos.**

(A) Heatmaps showing the expression of H3K4me2 related regulators in mouse (based on published RNA-seq data43).

(B) The box plots showing DNA methylation level at all promoter regions and distal region in mouse 2-cell stage. Random peaks are shown as a control.

(C-D) Venn diagrams showing De novo 4-cell H3K4me2 promoter and distal peaks number.

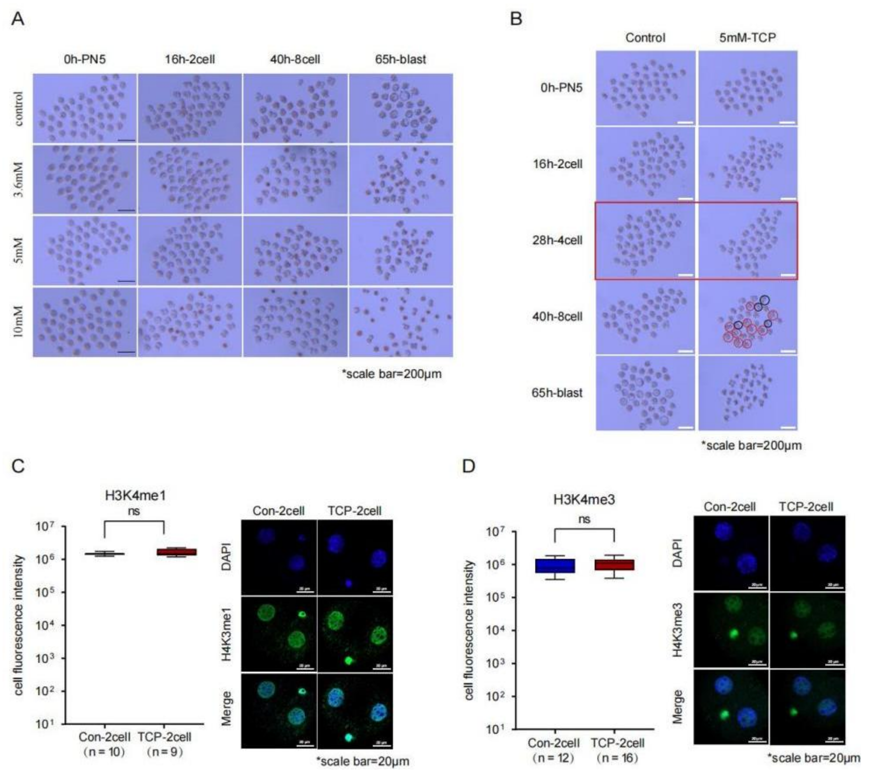


**Extended Data Figure 4. Cell culture experiment under TCP treatment**

(A)Morphological observation of cells under different concentration of TCP culture, scale bar is also shown.

(B)Morphological observation of embryo development after 5mM TCP inhibition of LSD2 expression.

(C-D)Immunofluorescence images showed the immunofluorescence intensity of H3K4me1 (C) and H3K4me3 (D) of 2-cell embryos in the experimental group (5mM TCP) and control group, scale bar is also shown.

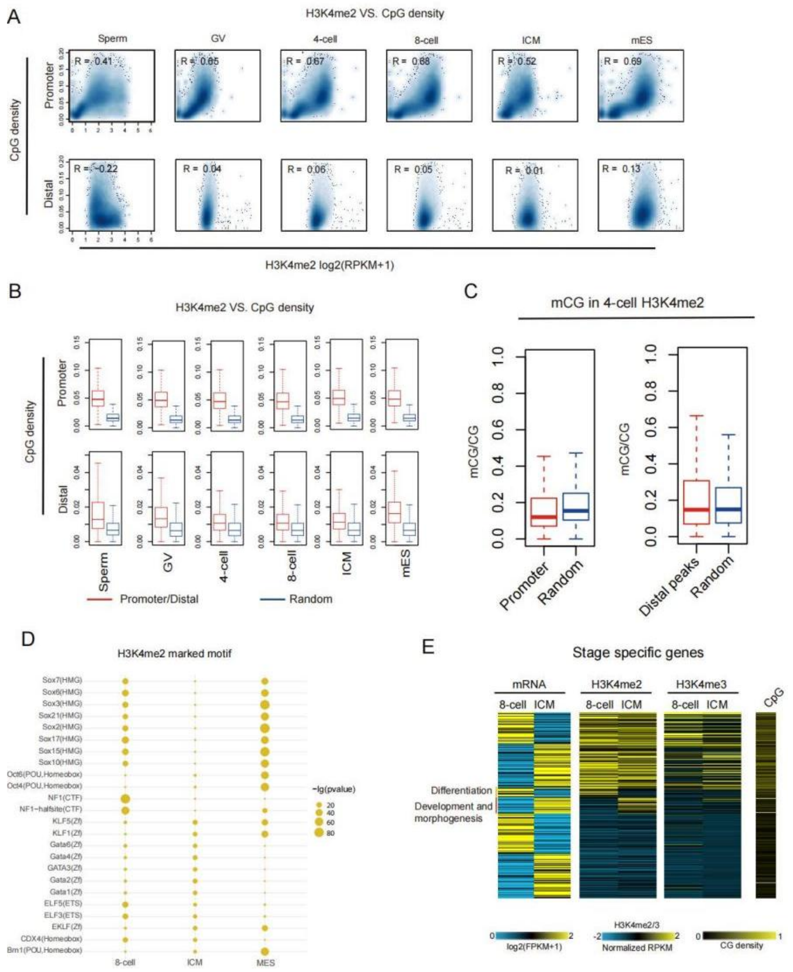


**Extended Data Figure 5. H3K4me2 in mouse 4/8-cell embryos.**

(A) Scatter plots comparing the promoter and distal H3K4me2 signals with CpG densities in mouse gametes, 4-cell, 8-cell, ICM and mESC. The Spearman correlation coefficients are also shown.

(B) The box plots showing the CpG density and promoter/distal H3K4me2 peaks. Random peaks are shown as a control.

(C) The box plot showing DNA methylation in promoter and distal 4-cell H3K4me2 peaks. Random peaks are shown as a control.

(D) TF motifs identified from active enhancers in mouse early embryos (8-cell, ICM) and mESC. showing motif enrichment p value < 1e-20 at least at one stage were included. Circle size showing TF enrichment and the expression of the TF is color coded.

(E) Heatmaps showing stage-specifically expressed genes further classified by their H3K4me2 states (high, dynamic, and low). RNA-seq, H3K4me3 state, CpG densities and GO analysis results are also mapped and shown.

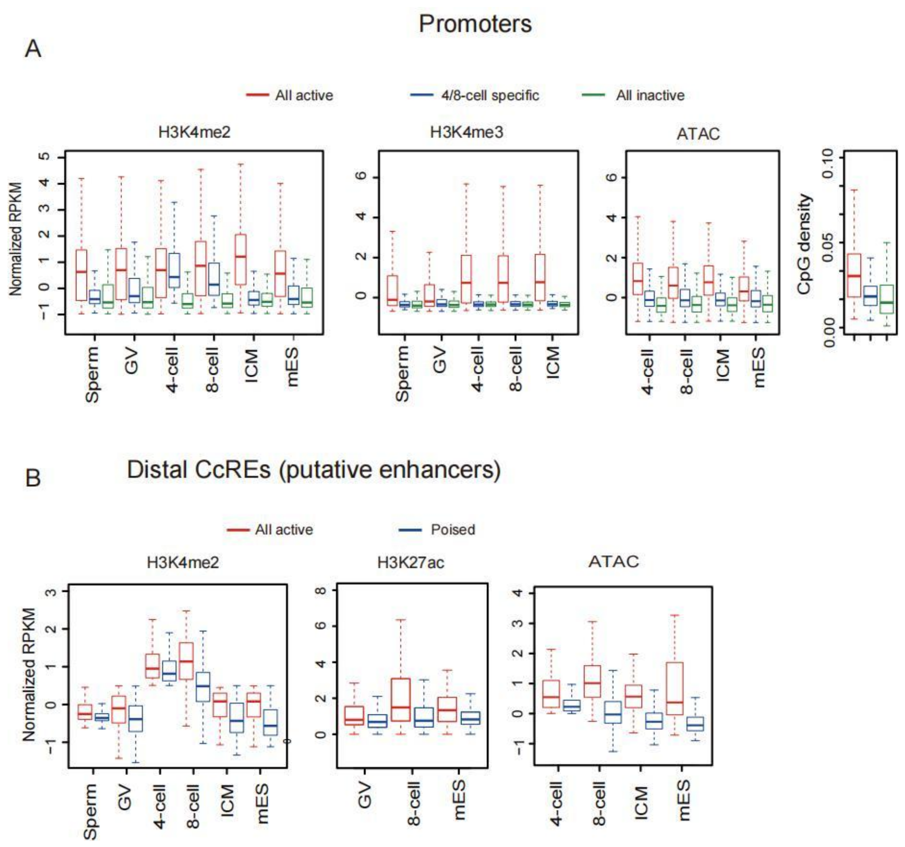


Extended Data Figure 6. Chromatin states at specific stages and distal CcREs.

(A) Box plots showing H3K4me2, H3K4me3 and ATAC-seq signals, “All active” and “All inactive” promoters in mouse gametes, embryos and mESC. CpG density for each group is also shown (right).

(B) The box plot showing the ATAC-seq, H3K4me2 and H3K27ac signals at putative active, poised or random enhancers in mouse gametes, early embryos or mESC, Mouse histone modified H3K27ac data refer to GEO database : GSE72784.

**Extended Data Table**

The sample information in this study was listed in the table below.

Table1. CUT&RUN experimental sample information

| Sample name | Experiment | Replicate | Sample number | Cell number |
|---|---|---|---|---|
| K562 cell line | H3K4me2CUT&RUN | Rep1 | 50 | 50 |
| K562 cell line | H3K4me2CUT&RUN | Rep2 | 50 | 50 |
| K562 cell line | H3K4me2CUT&RUN | Rep1 | 100 | 100 |
| K562 cell line | H3K4me2CUT&RUN | Rep2 | 100 | 100 |
| K562 cell line | H3K4me2CUT&RUN | Rep1 | 200 | 200 |
| K562 cell line | H3K4me2CUT&RUN | Rep2 | 200 | 200 |
| K562 cell line | H3K4me2CUT&RUN | Rep1 | 500 | 500 |
| K562 cell line | H3K4me2CUT&RUN | Rep2 | 500 | 500 |
| K562 cell line | H3K4me2CUT&RUN | Rep1 | 5000 | 5000 |
| K562 cell line | H3K4me2CUT&RUN | Rep2 | 5000 | 5000 |
| K562 cell line | H3K4me2CUT&RUN | Rep1 | 10000 | 10000 |
| K562 cell line | H3K4me2CUT&RUN | Rep2 | 10000 | 10000 |
| MES cell line | H3K4me2CUT&RUN | Rep1 | 50 | 50 |
| MES cell line | H3K4me2CUT&RUN | Rep2 | 50 | 50 |
| MES cell line | H3K4me2CUT&RUN | Rep1 | 100 | 100 |
| MES cell line | H3K4me2CUT&RUN | Rep2 | 100 | 100 |

| Sample name | Experiment | Replicate | Sample number | Cell number |
|---|---|---|---|---|
| MES cell line | H3K4me2CUT&RUN | Rep1 | 200 | 200 |
| MES cell line | H3K4me2CUT&RUN | Rep2 | 200 | 200 |
| MES cell line | H3K4me2CUT&RUN | Rep1 | 500 | 500 |
| MES cell line | H3K4me2CUT&RUN | Rep2 | 500 | 500 |
| MES cell line | H3K4me2CUT&RUN | Rep1 | 5000 | 5000 |
| MES cell line | H3K4me2CUT&RUN | Rep2 | 5000 | 5000 |
| MES cell line | H3K4me2CUT&RUN | Rep1 | 10000 | 10000 |
| MES cell line | H3K4me2CUT&RUN | Rep2 | 10000 | 10000 |
| GV oocyte | H3K4me2CUT&RUN | Rep1 | 190 | 190 |
| GV oocyte | H3K4me2CUT&RUN | Rep2 | 190 | 190 |
| MII oocyte | H3K4me2CUT&RUN | Rep1 | 165 | 165 |
| MII oocyte | H3K4me2CUT&RUN | Rep2 | 165 | 165 |
| PN5 zygote | H3K4me2CUT&RUN | Rep1 | 250 | 250 |
| PN5 zygote | H3K4me2CUT&RUN | Rep2 | 250 | 250 |
| Early_2cell | H3K4me2CUT&RUN | Rep1 | 86 | 172 |
| Early_2cell | H3K4me2CUT&RUN | Rep2 | 86 | 172 |
| Late_2cell | H3K4me2CUT&RUN | Rep1 | 98 | 196 |
| Late_2cell | H3K4me2CUT&RUN | Rep2 | 98 | 196 |
| 4cell | H3K4me2CUT&RUN | Rep1 | 65 | 260 |
| 4cell | H3K4me2CUT&RUN | Rep2 | 65 | 260 |

| Sample name | Experiment | Replicate | Sample number | Cell number |
|---|---|---|---|---|
| 8cell | H3K4me2CUT&RUN | Rep1 | 25 | 200 |
| 8cell | H3K4me2CUT&RUN | Rep2 | 25 | 200 |
| ICM | H3K4me2CUT&RUN | Rep1 | 14 | ≈420 |
| ICM | H3K4me2CUT&RUN | Rep2 | 14 | ≈420 |
